# *Pten* Orchestrates Neurogenic Radial Glia Lineage Progression and Tunes Neocortical Astrocyte Production

**DOI:** 10.64898/2026.05.01.722191

**Authors:** Osvaldo A. Miranda, Ximena Contreras, Florian M. Pauler, Amarbayasgalan Davaatseren, Nicole Amberg, Carmen Streicher, Ana Villalba, Anna Heger, Corentine Marie, Bassem A. Hassan, Thomas Rülicke, Simon Hippenmeyer

**Affiliations:** Institute of Science and Technology Austria, Am Campus 1, 3400 Klosterneuburg, Austria; Medical University of Vienna, Department of Neurology, Vienna, Austria; Institut du Cerveau-Paris Brain Institute (ICM), Sorbonne Universite, Inserm, CNRS, Hopital Pitie-Salpetriere, Paris, France; Yale School of Medicine, Section of Comparative Medicine, New Haven, CT 06520, USA; University of Veterinary Medicine Vienna, Institute of Laboratory Animal Science, Vienna, Austria; Medical University of Vienna, Ludwig Boltzmann Institute for Hematology and Oncology, Vienna, Austria

## Abstract

The cerebral cortex consists of immense numbers of neuronal and glial cell-types derived from radial glial progenitor (RGP) cells. How RGPs generate appropriate quantities of distinct cortical cell-types to safeguard a brain of correct size, is not well understood. However, genetic aberration in human, including mutations in *PTEN*, lead to cortical malformation such as macrocephaly, albeit with unknown etiology. Here we utilized Mosaic Analysis with Double Markers (MADM)-based clonal analysis and single cell phenotyping to decipher the role of *Pten* in neurogenic and gliogenic RGP lineage progression during cortical ontogeny. While neurogenic RGP lineage progression and projection neuron production was moderately altered in the absence of *Pten*, cortical astrocyte production was drastically increased. Through genetic epistasis experiments we show that the loss of *Pten* uncouples astrocyte generation from essential growth factor signaling hubs, funneling into MAPK. Collectively, our results suggest that *Pten* regulates RGP lineage progression with distinct sequential functions in cortical projection neurogenesis and astrocyte production to ensure the emergence of a correctly-sized cerebral cortex.

## INTRODUCTION

The cerebral cortex is the entity that instructs higher order cognitive brain functions. The mature cerebral cortex consists of a diverse population of neuronal and glial cell types that organize into six distinct cytoarchitectural laminae (Molnar et al. 2019; Hanganu-Opatz et al. 2021; Namba and Huttner 2024; Nowakowski et al. 2025). During cerebral cortex ontogeny, precise developmental programs instruct neural stem cells (NSCs) to generate the correct range of neuronal and glial cell types and thereby ensure the establishment of correctly sized brains. Impairments in cortical neurogenesis and/or glia production lead to alterations in the cortical cytoarchitecture, which is thought to reflect the underlying cause for neurodevelopmental disorders, including cortical malformations such as microcephaly or megalencephaly, autism, intellectual disability and epilepsy (Barkovich et al. 2012; Pirozzi et al. 2018; Sullivan and Geschwind 2019; Klingler et al. 2021; Bizzotto and Walsh 2022; Asif et al. 2023).

During embryogenesis, the early neuroepithelium is composed of neuroepithelial stem cells (NESCs) that initially amplify their pools but eventually transform into radial glial progenitor (RGPs) cells (Kriegstein and Alvarez-Buylla 2009; Taverna et al. 2014; Villalba et al. 2021). RGPs are the major progenitor cells from which all cortical projection neuron lineages and certain macroglial populations derive. The proliferation behavior and temporal lineage progression of neurogenic and gliogenic RGPs needs to be tightly regulated but the underlying cellular and molecular mechanisms at the individual cell level are not well understood.

While major research in the past has provided a rough framework of cortical neurogenesis and glia production (Freeman and Rowitch 2013; Greig et al. 2013; Casingal et al. 2022; Clavreul et al. 2022; Lattke and Guillemot 2022; Bartels et al. 2024; Di Bella et al. 2024), recent single cell lineage tracing experiments in mice, utilizing Mosaic Analysis with Double Markers (MADM) technology (Zong et al. 2005; Contreras et al. 2021), has revealed that cortical RGP lineage progression exhibits a high level of stereotypy with predictable quantitative parameters (Lin et al. 2021; Llorca and Marin 2021; Hippenmeyer 2023). As such, RGPs initially divide symmetrically to amplify their numbers before they switch to asymmetric neurogenic division mode at around embryonic day (E) 12 (Gao et al. 2014). During the phase of asymmetric neurogenic division, RGPs produce a predictable unit of ∼8-9 projection neurons, either directly or indirectly via intermediate progenitors (IPs) (Gao et al. 2014; Llorca et al. 2019). Once neurogenesis is completed, certain subpopulations of RGPs give rise to ependymal cells, transform into type B1 cells located in the ventricular-subventricular zone (V-SVZ) (Beattie et al. 2017; Ortiz-Alvarez et al. 2019), or transform into gliogenic progenitors that establish astrocyte and/or oligodendrocyte lineages in the developing cortex (Gao et al. 2014; Zhang et al. 2020; Shen et al. 2021). At the quantitative level, about one in six RGPs adopt glial potential whereby defined fractions of these gliogenic RGPs produce astrocyte-or oligodendrocyte intermediate progenitors (aIPs or oIPs) and their subsequent glial lineages (Gao et al. 2014; Beattie et al. 2017; Zhang et al. 2020; Shen et al. 2021). In contrast to neurogenic RGPs, with their cellular attachments at the ventricular and pial surfaces (Taverna et al. 2014; Villalba et al. 2021), gliogenic progenitors locate throughout the cortical wall and individual aIPs generate astrocyte lineages with high numeric variance (Ge et al. 2012; Gao et al. 2014; Clavreul et al. 2019; Zhang et al. 2020; Shen et al. 2021). The current model of astrocyte genesis postulates that due to the characteristic tiling organization of mature astrocytes, aIPs follow a space-filling mode of proliferation (Clavreul et al. 2022; Markey et al. 2023; Bartels et al. 2024; Chung et al. 2024; Williamson et al. 2025). Although the above framework of neurogenic and gliogenic RGP lineage progression provides clear temporal operational hubs and predictable quantitative parameters, the cellular and molecular mechanism regulating the sequential steps of lineage progression and RGP neuron and glia output potential remain mostly unknown.

Here we focused on the role of phosphatase and tensin homologue on chromosome ten (PTEN). In human, deleterious mutations in *PTEN* are associated with a variety of neurodevelopmental disorders but most prominently with cortical malformations such as macrocephaly (i.e. bigger brain), hemimegalencephaly, and focal cortical dysplasia (Skelton et al. 2020; Ma et al. 2024; Currey et al. 2025; Fazekas et al. 2026). Conversely, a microduplication of 10q21.31 affecting three genes, including *PTEN* as the putative causal gene, has been identified in patients with autosomal dominant primary microcephaly (Oliveira et al. 2019). Hence, *PTEN* appears to represent a major signaling hub for controlling brain size in human. At the biochemical level, PTEN antagonizes the Phosphatidyl Inositide 3-Kinase (PI3K)-AKT-mammalian Target of Rapamycin Complex 1 (mTORC1) pathway, and thereby regulates a myriad of signaling transduction pathways (Song et al. 2012; Veleva-Rotse and Barnes 2014; Worby and Dixon 2014). Previous loss of function (LOF) studies in mice have revealed critical *Pten* functions during cortical development albeit mostly at the macroscopic level. While complete knockout of *Pten* results in early embryonic lethality (Di Cristofano et al. 1998), conditional whole brain (Backman et al. 2001; Groszer et al. 2001; Kwon et al. 2001), or cortex-specific (Lehtinen et al. 2011; Chen et al. 2015) elimination of *Pten* led to macrocephaly, resembling to some extent the observed clinical features associated with *PTEN* mutation in human. Interestingly, conditional elimination of *Pten* in astrocyte lineages also showed progressive enlargement of the brain (Fraser et al. 2004). Most of the above research clearly showed a larger cerebral cortex / brain upon loss of *Pten*, and therefore imply a critical role of *Pten* in cortical neurogenesis and/or astrocyte development. However, the functional cell-autonomous requirement of *Pten* in neurogenic and gliogenic RGP lineage progression at single progenitor cell level in the context of the current framework is not known. We therefore capitalized upon the high resolution MADM technology, enabling single cell genetics and RNA sequencing in combination with lineage tracing, to probe the role of *Pten* quantitatively and qualitatively with the goal to obtain definitive insight at the individual RGP cell level.

## RESULTS

### Loss of *Pten* affects temporal neurogenic RGP lineage progression and results in enlarged RGP-derived unitary projection neuron output

The ablation of *Pten* function during development in mouse results in macrocephaly and enlarged cerebral cortex (Figure 1A). How the loss of *Pten* phenotypically translates at the single cell level and, in particular, how RGP lineage progression and neuron production may be affected is not clear (Figure 1B). To this end, we conceived of two genetic MADM-based paradigms enabling quantitative phenotypic assessment and therefore evaluation of cell-autonomous *Pten* function in RGP-mediated neurogenesis with true single cell resolution. First, we generated a sparse genetic mosaic with *Pten* deletion in just very few *Emx1^+^* RGPs and their corresponding projection neuron lineages (*Pten*-MADM, *MADM-19^GT/TG,Pten^*;*Emx1^Cre/+^*; see also Methods and Figure S1 for details). In mosaic *Pten*-MADM, red tdT^+^ MADM-labeled cells were *Pten^+/+^* (wild-type) and green GFP^+^ cells *Pten^-/-^* (homozygous mutant). To normalize our data set, we compared *Pten*-MADM to Control-MADM (*MADM-19^GT/TG^*;*Emx1^Cre/+^*) where red tdT^+^ and green GFP^+^ MADM-labeled cells were *Pten^+/+^*(Figure S1). As expected, the numbers of green (G) and red (R) projection neurons in Control-MADM at postnatal day (P) 21 were identical and we thus calculated a G/R ratio of ∼1 (Figure 1C-1E). In contrast, the G/R ratio in *Pten*-MADM was ∼2 with about twice as many green *Pten^-/-^*projection neurons when compared to red *Pten^+/+^* control cells (Figure 1F-1H). Next, we assessed G/R ratios in Control- and *Pten*-MADM during embryonic development to identify the critical temporal window of phenotype emergence (Figure S2). We quantified G/R ratios of PAX6^+^ progenitor cells and PAX6^-^ cells (IPs and nascent projection neurons) separately. At E12 (onset of neurogenesis), the numbers of both PAX6^+^ and PAX6^-^ cell populations were identical in *Pten*- and Control-MADM, respectively. While *Pten^-/-^* PAX6^+^ progenitor cells were significantly increased from E14 onward, PAX6^-^ cells showed a trend towards increase at E14 but were significantly increased at E16 (Figure S2O-S2P). Thus, the homozygous loss of *Pten* results in an increase of PAX6^+^ RGPs and their corresponding nascent IP/projection neuron lineages during embryonic development.

**Figure 1.**
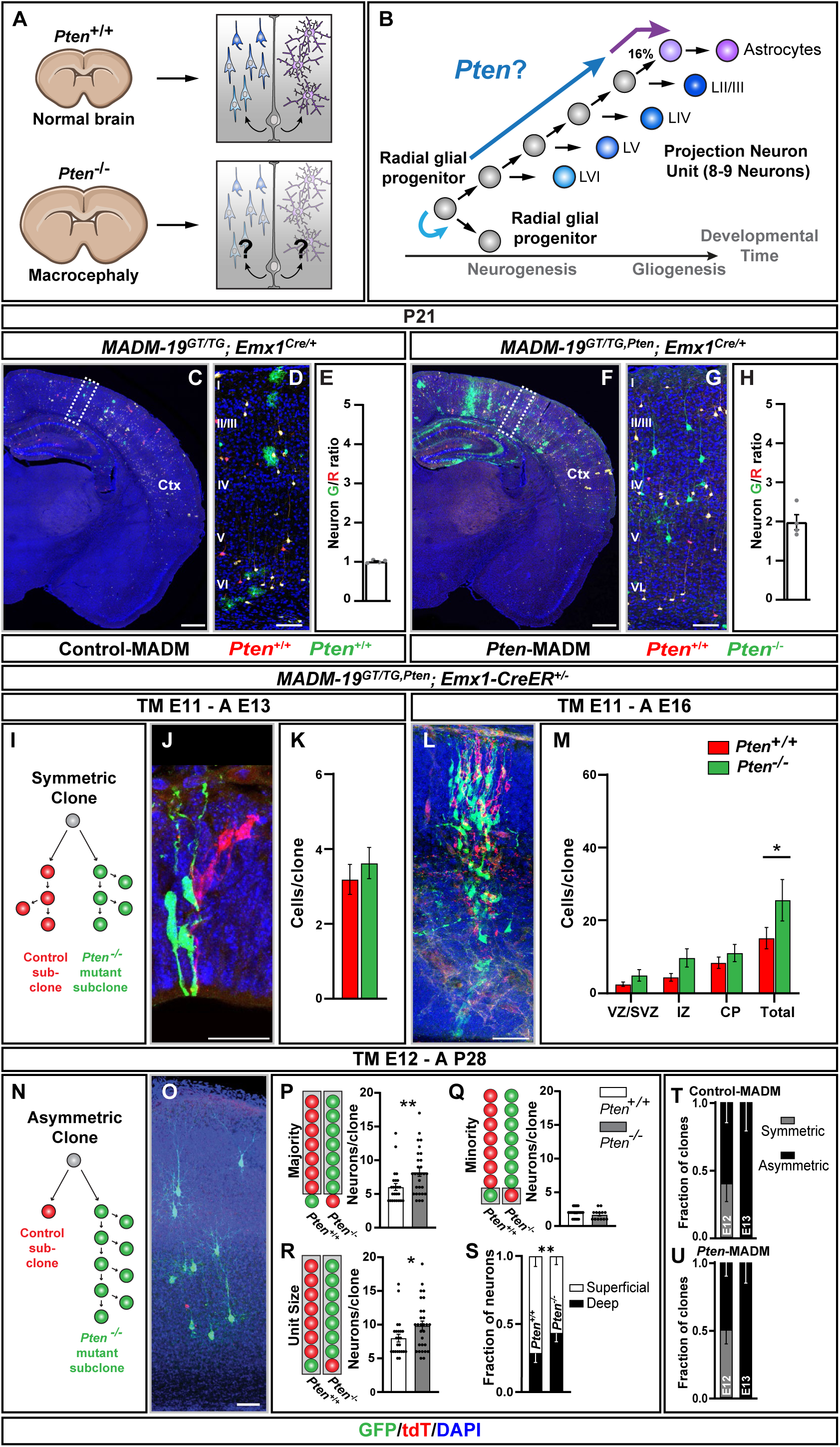
*Pten* cell-autonomously controls RGP-mediated neuron output during corticogenesis. (A) *Pten* loss of function (LOF) results in macrocephaly with enlarged cerebral cortex, but the underlying basis of the phenotype at single cell level is unclear. (B) Quantitative framework of radial glial progenitor (RGP) cell lineage progression in the developing cortex with putative role(s) of *Pten* in regulating critical stages and transitions. (**C-H**) Representative images of somatosensory cortex in overview (C and F), higher resolution of cortical plate (D and G), and quantification of G/R projection neuron ratio in *MADM-19^GT/TG^*;*Emx1^Cre/+^* (C-E; Control-MADM with red tdT^+^ *Pten^+/+^* cells and green GFP^+^ *Pten^+/+^* cells) and *MADM-19^GT/TG,Pten^*;*Emx1^Cre/+^* (F-H; *Pten*-MADM with red tdT^+^ *Pten^+/+^* cells and green GFP^+^ *Pten^-/-^* cells). Note that G/R ratio is increased in *Pten*-MADM, indicating a larger GFP^+^ *Pten^-/-^* mutant neuron population when compared to tdT^+^ control neuron numbers. (**I-M**) MADM clonal analysis and quantification of neuron output in symmetric MADM clones. (I) Schema of a symmetric MADM clone, with a control tdT^+^ *Pten^+/+^* subclone and GFP^+^ *Pten^-/-^* mutant subclone. (J-M) Representative images of MADM clones (J and L), and quantification of MADM-labeled cells (K and M) in *MADM-19^GT/TG,Pten^*;*Emx1-CreER^+/-^* with MADM clones induced at E11 and analyzed at E13 (J) or E16 (L). (**N-R**) MADM clonal analysis and quantification of neuron output in asymmetric MADM clones. (N) Schema of an asymmetric MADM clone, with a control tdT^+^ *Pten^+/+^* subclone and GFP^+^ *Pten^-/-^* mutant subclone. (O-R) Representative image of an asymmetric MADM clone (O), and quantification of majority (P), minority (Q), and total unit size (R) in *MADM-19^GT/TG,Pten^*;*Emx1-CreER^+/-^* with MADM clones induced at E12 and analyzed at P28. (**S**) Quantification of cortical projection neuron fractions across deep (VI-V) and superficial (IV-II) layers in *MADM-19^GT/TG^*;*Emx1-CreER^+/-^*and *MADM19^GT/TG,Pten^*;*Emx1-CreER^+/-^*. (**T-U**) The fraction of MADM clones in either symmetric proliferative division or asymmetric neurogenic division at the time of induction in *MADM-19^GT/TG^*;*Emx1-CreER^+/-^* and *MADM-19^GT/TG,Pten^*;*Emx1CreER^+/-^*. Nuclei were stained using DAPI (blue). Ctx: Neocortex. Cortical layers are indicated in roman numerals. Values represent mean ± SEM. Significance was determined using two-way ANOVA with Šídák’s multiple comparison test (M), Mann-Whitney test (P-R), and Fischer’s exact test (S). *p < 0.05, **p < 0.01. Scale bar, 500 μm (C and D), 50 μm (D, G and O), 25 μm (J and L).

To identify the critical *Pten*-dependent stage(s) in neurogenic RGP lineage progression, and to monitor RGP-mediated unitary projection neuron output upon *Pten* loss of function at the individual RGP level, we next conducted MADM-based clonal analysis (Gao et al. 2014; Beattie et al. 2020). To induce MADM clones we utilized the temporally-inducible tamoxifen (TM)-dependent *Emx1*-CreER driver (Kessaris et al. 2006). First, we analyzed RGPs in their symmetric proliferation mode where the two nascent MADM-labeled daughter cells and corresponding subclones, emerging from a dividing RGP, will be labeled in red (tdT^+^) or green (GFP^+^) fluorescent colors. To this end, we induced MADM clones at E11 in *MADM-19^GT/TG,Pten^*;*Emx1-CreER^+/-^*; when a majority of RGPs undergo symmetric proliferative divisions (Gao et al. 2014). In such a scenario, the red tdT^+^ MADM subclone consists of *Pten^+/+^*cells whereas the green GFP^+^ subclone was *Pten^-/-^* (Figure 1I). We next analyzed the sizes of the respective subclones at E13 and E16 (end of neurogenic period) and found a trend towards increased numbers of *Pten^-/-^* mutant cells that got significant over the total clone size at E16 (Figure 1I-1M), which is in agreement with the above data derived from sparse genetic mosaic.

In the next experiment, we assessed projection neuron output of single RGPs upon their switch to asymmetric neurogenic division mode (Gao et al. 2014; Beattie et al. 2020). We induced asymmetric MADM clones at E12 and analyzed projection neuron numbers at P28. In such a setting, the red tdT^+^ subclone again consisted of *Pten^+/+^* cells whereas the green GFP^+^ subclone contained *Pten^-/-^* cells (Figure 1N). We quantified total clone/unit size, majority population (larger subclone) and minority population (smaller subclone) (Figure 1O-1R). While there was no difference in the sizes of the minority subclones (Figure 1Q), the number of *Pten^-/-^*cells in majority population and overall unit size was increased in the absence of *Pten* (Figure 1P and 1R). Interestingly, the relative fraction of deep layer V and VI neurons was increased in *Pten^-/-^* mutant clones when compared to control clones (Figure 1S), a finding that was corroborated by analyzing genetic *Pten*-MADM mosaics (Figure S3). We also noticed a significant increase of cell size in *Pten^-/-^*mutant neurons (Figure S4), in agreement with previous studies (Groszer et al. 2001; Fraser et al. 2008). Next, we quantified the relative proportions of symmetric proliferative versus asymmetric neurogenic MADM clones when induced at E12 and E13 with analysis at P28 (Figure 1T and 1U). While a majority of Control-MADM clones (i.e. red and green cells were *Pten^+/+^*) had transitioned to asymmetric division when induced at E12 (less than 37% symmetric division), a larger fraction (∼50%) of clones in *Pten*-MADM appeared symmetric. However, all MADM clones appeared in asymmetric architecture when induced at E13 (Figure 1T and 1U). These data imply that symmetric proliferative RGP division persisted slightly longer in the absence of *Pten*, which is in agreement with the observation of increased overall numbers of *Pten^-/-^* RGPs (Figure S2). In summary, genetic loss of *Pten* in cortical RGPs leads to an extension of symmetric RGP division mode and thereby higher numbers of RGPs, and a slight but significant increase in the RGP-derived overall projection neuron unit size. Collectively, due to aberrant neurogenic RGP lineage progression and increased neuron output *Pten^-/-^* mutant cortical projection neurons were significantly increased.

### Loss of *Pten* leads to increased numbers of cortical astrocytes

Upon completion of neurogenesis, a fraction of cortical RGPs attain gliogenic potential and proceed with the generation of astrocytes (Gao et al. 2014; Beattie et al. 2017; Zhang et al. 2020; Shen et al. 2021). While previous studies, utilizing conditional cortex-wide deletion of *Pten*, have shown hypertrophy and increased proliferation of cortical astrocytes (Fraser et al. 2004), the underlying mechanisms are unknown. To quantitatively approach these issues, we first analyzed the sparse genetic MADM mosaics at P21. While in Control-MADM, green and red astrocyte numbers were both ∼1 (Figure 2A-C), we noticed that *Pten^-/-^*mutant astrocytes were increased by about five-fold in *Pten*-MADM (Figure 2D-2F). In agreement with literature (Fraser et al. 2004), *Pten^-/-^* mutant astrocytes also showed increased nuclear and cellular size, and significantly more complex branching patterns (Figure S5).

**Figure 2.**
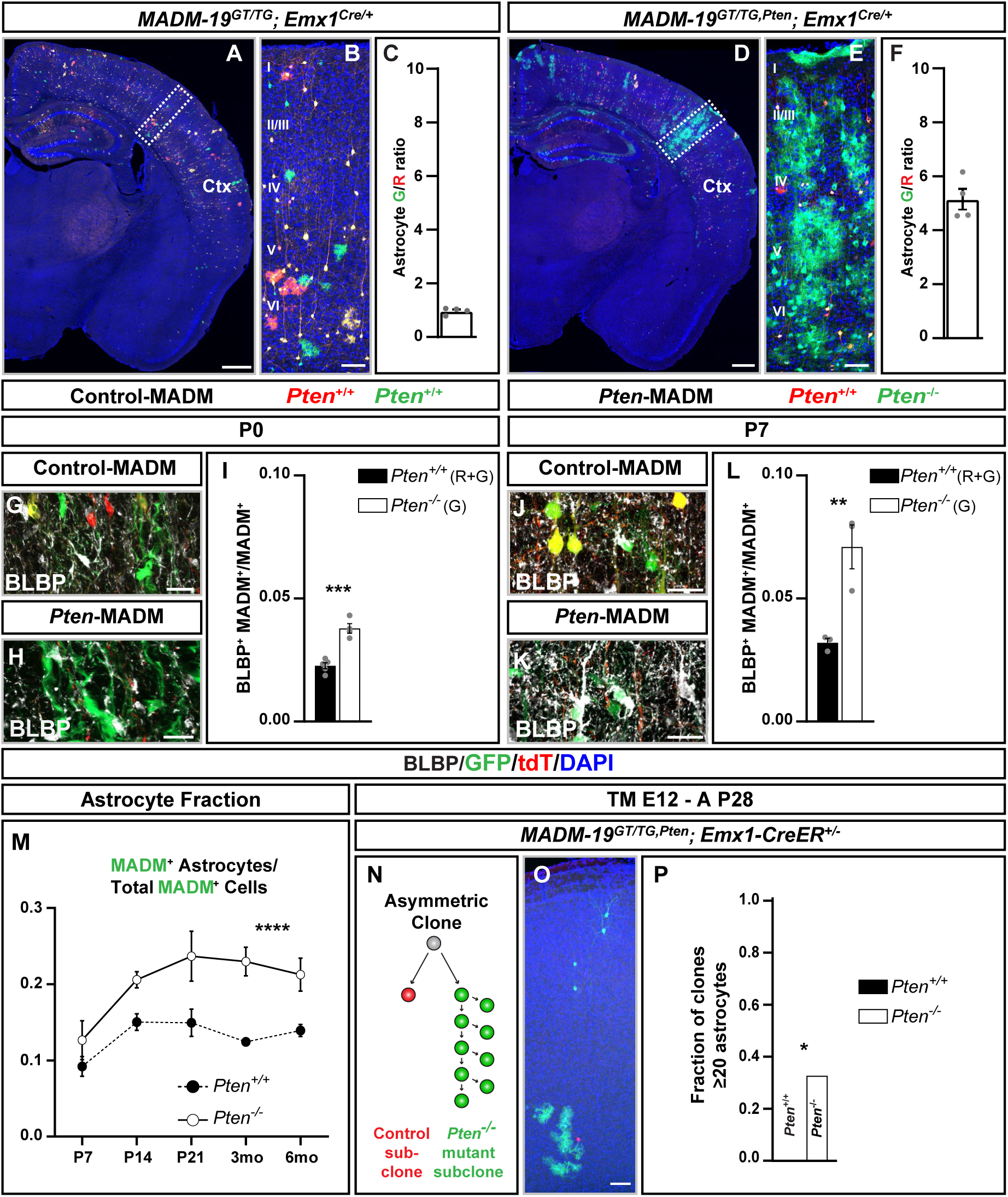
*Pten* loss of function leads to increased cortical astrocyte production. (**A-F**) Representative images of somatosensory cortex in overview (A and D), higher resolution of cortical plate (B and E), and quantification of G/R astrocyte ratio in *MADM-19^GT/TG^*;*Emx1^Cre/+^* (A-C; Control-MADM with red tdT^+^ *Pten^+/+^* cells and green GFP^+^ *Pten^+/+^* cells) and *MADM-19^GT/TG,Pten^*;*Emx1^Cre/+^* (D-F; *Pten*-MADM with red tdT^+^ *Pten^+/+^* cells and green GFP^+^ *Pten^-/-^* cells). Note that G/R ratio is increased in *Pten*-MADM, indicating a larger GFP^+^ *Pten^-/-^* mutant astrocyte population when compared to tdT^+^ control numbers. (**G-L**) Representative images of BLBP^+^ aIPs in cortical plate (G, H, J, K), and quantification of the fraction of BLBP^+^GFP^+^ cells over total GFP^+^ population (I and L) in Control-MADM (G, I, J and L) and *Pten*-MADM (H, I, K and L) at P0 (G-I) and P7 (J-L). (**M**) Time course of fraction of GFP^+^ astrocytes in Control-MADM (white) and *Pten*-MADM (black) at P7, P14, P21, 3 months, and 6 months of age. (**N-P**) MADM clonal analysis and quantification of astrocyte output in asymmetric MADM clones in *MADM-19^GT/TG,Pten^*;*Emx1CreER^+/-^*, induced at E12 and analyzed at P28. (N) Schema of an asymmetric MADM clone, with a control tdT^+^ control subclone and GFP^+^ *Pten^-/-^* mutant subclone. (O-P) Representative image of an asymmetric MADM clone (O), and quantification of the fraction of large astrocyte clones (greater than or equal to 20 astrocytes) in control and *Pten^-/-^* mutant subclones (P). Note that large astrocyte clones were never observed in control subclones but that more than thirty percent of *Pten* mutant asymmetric clones consisted of large astrocyte clones. Nuclei were stained using DAPI (blue). Ctx: Neocortex. Cortical layers are indicated in roman numerals. Values represent mean ± SEM. Significance was determined using unpaired two-tailed t-test (I and L), two-way ANOVA with Šídák’s multiple comparison test (M), and Fischer’s exact test (P). *p < 0.05, ***p < 0.001, ****p < 0.0001. Scale bar, 500 μm (A and D), 50 μm (B, E and O), 25 μm (G, H, J and K)

Next, we assessed the origin of increased *Pten^-/-^* mutant astrocyte numbers in *Pten*-MADM and quantified the number of BLBP^+^ aIPs. At P0 and P7 the numbers of *Pten^-/-^*mutant aIPs were significantly increased in *Pten*-MADM when compared to Control-MADM (Figure 2G-2L). Thus, the loss of *Pten* in gliogenic RGPs leads to increased aIP production which subsequently results in higher numbers of cortical astrocytes. Furthermore, time course analysis revealed that astrocyte generation in Control-MADM plateaued after the second postnatal week while in *Pten*-MADM, increased cortical astrocyte production continued up to the third postnatal week (Figure 2M). *Pten^-/-^* mutant astrocytes were often found in columnar arrangement and close proximity to each other. We therefore next generated individual MADM clones to determine the clonal astrocyte output from single RGPs. We induced MADM clones at E12 to obtain primarily asymmetric RGP-derived clonal units (Figure 2N) and observed a significant fraction of *Pten^-/-^* mutant MADM clones containing ≥20 astrocytes, whereas *Pten^+/+^* clones never contained >20 astrocytes in our experimental paradigm (Figure 2P). Altogether, the loss of *Pten* in gliogenic RGPs leads to significantly elevated numbers of cortical astrocytes due to increased generation of aIPs and overall higher clonal astrocyte output during a prolonged gliogenic period.

### *Pten^-/-^* mutant astrocytes show deregulation of genes associated with key growth factor signaling pathways

In an effort to obtain deeper insights into the molecular correlates associated with the loss of *Pten* function phenotype in cortical astrocyte production, we first conceived a bulk RNA sequencing strategy (Figure 3A). With the goal to obtain ultrapure astrocyte lineages for sequencing (Laukoter et al. 2020b; Amberg et al. 2022), we crossed *Pten*-MADM animals to mice carrying a transgenic *lacZ* reporter [*hGFAP-lacZ*; (Brenner et al. 1994)], specifically expressed in cortical astrocytes (Figure 3A). MADM-labeled cells were then sorted from Control-MADM (with GFP^+^/lacZ^+^; *Pten^+/+^*cells) and *Pten*-MADM (with GFP^+^/lacZ^+^; *Pten^-/-^*cells) at P0 and P4, and subjected to RNA sequencing (see Methods for details). For subsequent bioinformatics analysis we first carried out principal component analysis (PCA) using the top 500 variable genes and observed that experimental samples clustered according to age and genotype (Figure S6A). Next, we validated the deletion of *Pten* exon 5 in cells originated from *Pten*-MADM (Figure S6B). We assessed differentially expressed genes (DEGs) and noticed an increased number of DEGs at P0 when compared to P4 (Figure 3B), and that those DEGs were distinct at each time point (Figure 3C). By using gene ontology (GO) assignment, we found that a large number of DEGs were associated with gliogenesis. We also found a number of DEGs implicated with key growth factor signaling such as genes encoding for FGFR and NOTCH receptors as well as *Etv5* (Figure 3D). To more systematically analyze gene regulatory networks (GRNs) associated with *Pten* knockout we pursued STRING [Search Tool for the Retrieval of Interacting Genes/Proteins, (Szklarczyk et al. 2023)] analysis. We included selected key genes that have been associated previously with cortical astrocyte genesis (Figure 3E) in order to evaluate if and how they integrate into the GRN. We found a major hub associated with *Mki67* and cell-cycle regulation at P0, but also prominent clusters around genes related to growth factor signaling, including *Egfr*, *Notch1*, *Jak1/2*, *Mapk1/3* at both P0 and P4 stages (Figure 3F). In summary, the bulk transcriptomic data set suggests an acute deregulation of genes encoding for key cell cycle components and major growth factor signaling pathways.

**Figure 3.**
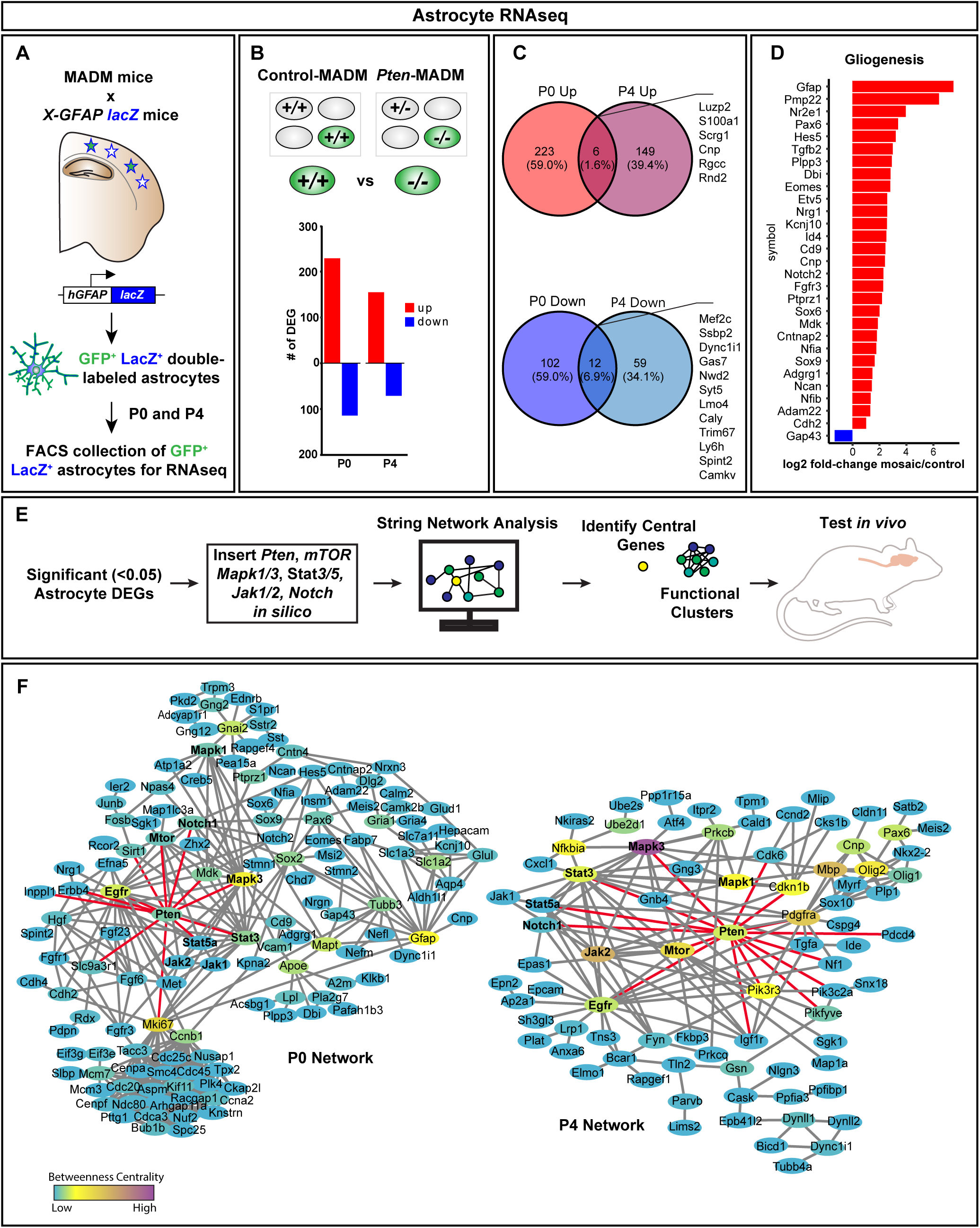
RNA-sequencing of highly pure astrocyte populations reveals deregulation of key signaling pathways in *Pten^-/-^* mutant astrocytes. (A) Experimental paradigm for the generation and FACS-based isolation of highly pure astrocyte populations. MADM mice were crossed with mice carrying a *lacZ* transgene under the control of the *hGFAP* promotor to generate GFP^+^LacZ^+^ double-positive astrocytes. Samples were collected at P0 and P4 and GFP^+^LacZ^+^ double-positive astrocytes subjected to RNA-seq. (B) Number of DEGs in GFP^+^LacZ^+^ astrocytes by comparison of Control-MADM and *Pten*-MADM (P_adj_<0.01, DESeq2) at P0 and P4. (C) Overlap of DEGs at P0 and P4 is indicated. Note that up- and downregulated DEGs display minimal overlap between P0 and P4. (D) Bar plot showing all up- and down-regulated genes associated with gliogenesis at P0. (E) Schematic of experimental workflow for STRING network analysis as hypothesis generator for identifying candidate genes for subsequent analysis *in vivo*. (F) STRING network analysis at P0 (left) shows a prominent clustering of cell cycle genes with *Mki67* as a highly central node in the network. Genes added *in silico* are marked with bold font. Red edges indicate first-degree neighbors of *Pten*. *Mapk3*, *Gfap*, and *Egfr* are central nodes in the network and, though these genes were added *in silico*, these data indicate a deregulation of genes associated with those pathways. The P4 STRING network (right) differs from the P0 network and many of the genes observed in the P0 network are not present at P4, notably the cluster of cell-cycle related genes is absent at P4.

### *Pten* deletion overrides the essential EGFR requirement for cortical astrocyte production

In order to assess the *in vivo* relevance of the above predicted DEGs and putative role in the *Pten* loss of function astrocyte phenotype, we pursued MADM-based epistasis experiments with selected candidate genes. First we focused on *Egfr* since previous studies have demonstrated an absolutely essential requirement for EGFR in the generation of cortical astrocytes (Burrows et al. 1997; Sibilia et al. 1998; Beattie et al. 2017; Zhang et al. 2020). We thus reasoned that the increased production of cortical astrocytes in the *Pten^-/-^* condition should abolished by eliminating *Egfr* and *Pten* at the same time, if the *Pten^-/-^*astrocyte phenotype was strictly dependent on *Egfr* (Figure 4A). To test our hypothesis, we generated the following genetic paradigm: *Pten*-MADM in *Egfr*-cKO (*Pten*-MADM-*Egfr*-cKO; *MADM-19^GT/TG,Pten^*;*Egfr^flox/flox^*;*Emx1^Cre/+^*) and assessed astrocyte production in comparison to Control-MADM (*MADM-19^GT/TG^*;*Emx1^Cre/+^*), *Pten*-MADM (*MADM-19^GT/TG,Pten^*;*Emx1^Cre/+^*) and Control-MADM-*Egfr*-cKO (*MADM-19^GT/TG^*;*Egfr^flox/flox^*;*Emx1^Cre/+^*) (Figure S7). To simplify the quantification, we assessed the absolute number of green MADM-labeled cortical astrocytes per mm^2^ (Figure 4B-4J). In Control-MADM (GFP^+^ cells were *Pten^+/+^*;*Egfr^+/+^*) we observed ∼5 astrocytes/mm^2^ (black bar in Figure 4J) whereas the number of GFP^+^ *Pten^-/-^*;*Egfr^+/+^*astrocytes was significantly increased in *Pten*-MADM (white bar in Figure 4J), in agreement with our above data (Figure 2). As expected and in line with our previous data (Beattie et al. 2017), in Control-MADM-*Egfr*-cKO (*MADM-19^GT/TG^*;*Egfr^flox/flox^*;*Emx1^Cre/+^*; green cells were GFP^+^ *Pten^+/+^*;*Egfr^-/-^*) we detected nearly no astrocytes (orange bar in Figure 4J). In contrast, concomitant deletion of *Pten* and *Egfr* in *Pten*-MADM-*Egfr*-cKO did result in a significant increase in GFP^+^ *Pten^-/-^*;*Egfr^-/-^* cortical astrocytes (blue bar in Figure 4J), comparable (i.e. no significant difference) to *Pten*-MADM (Figure 4J). Thus, the deletion of *Pten* appears to render cortical astrocyte production insensitive to EGFR status.

**Figure 4.**
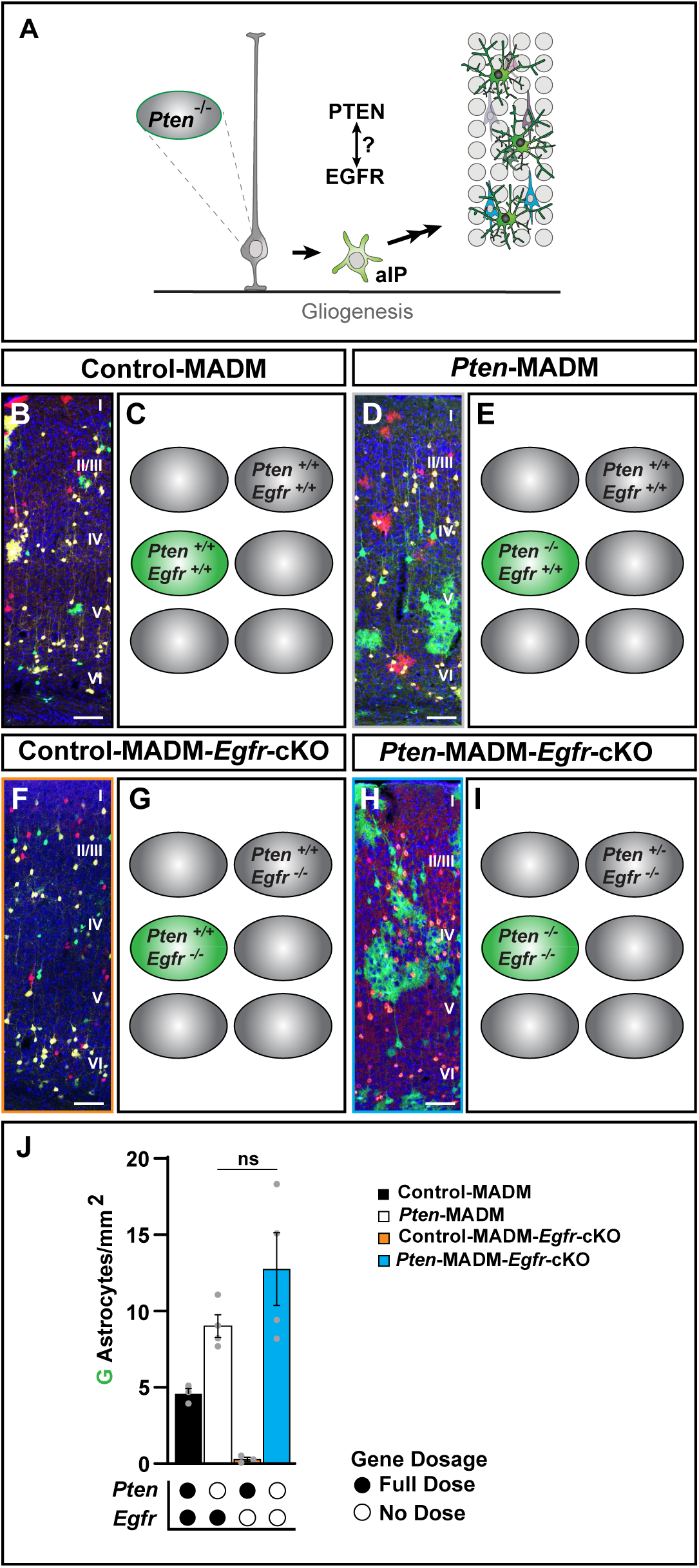
*Pten* loss of function overrides the essential *Egfr* requirement for cortical astrocyte production. **(A)** Schematic illustrating putative epistatic interaction between *Pten* and *Egfr* requirement for the production of cortical astrocytes. The concept of the experiment relied on the question whether the *Pten^-/-^*astrocyte overproduction phenotype could be rescued by concomitant deletion of *Egfr*. **(B-I)** Representative images of P21 somatosensory cortex (B, D, F, and H), and schematics illustrating the genotype in MADM-labeled GFP^+^ cells and the unlabeled background (C, E, G and I) in Control-MADM (B, C; *MADM-19^GT/TG^*;*Emx1^Cre/+^*), *Pten*-MADM (D, E; *MADM-19^GT/TG,Pten^*;*Emx1^Cre/+^*), Control-MADM-*Egfr*-cKO (F, G; *MADM-19^GT/TG^*;*Egfr^flox/flox^*;*Emx1^Cre/+^*), and *Pten*-MADM-*Egfr*-cKO (H, I; *MADM-19^GT/TG,Pten^;Egfr^flox/flox^*;*Emx1^Cre/+^*). (**J**) Quantification of astrocyte/mm^2^ in the P21 somatosensory cortex in Control-MADM (black), *Pten*-MADM (white), Control-MADM-*Egfr*-cKO (orange), and *Pten*-MADM-*Egfr*-cKO (blue). *Pten* and *Egfr* gene dosage is indicated below plot. Nuclei were stained using DAPI (blue). Cortical layers are indicated in roman numerals. Values represent mean ± SEM. Significance was determined using unpaired t-test. ns, non-significant. Scale bar, 50 μm (B, D, F and H).

### Single cell RNA sequencing reveals highly cell-type specific deregulation of gene expression upon loss of *Pten* and *Egfr*

To assess molecular underpinnings of *Egfr*-insensitivity upon *Pten* loss of function in cortical astrocyte generation we capitalized on single cell RNA sequencing (Figure 5). Specifically, we utilized particle-templated instant partition sequencing [PIP-seq; (Clark et al. 2023)] since this approach is tailored for lower-input sample sizes and ideally suited for MADM-labeled sparse cell populations. We isolated cortical tissue and processed single cell suspensions from Control-MADM, *Pten*-MADM and *Pten*-MADM-*Egfr*-cKO animals at P0. Next, we used FACS to isolate GFP^+^ cells from Control-MADM (*Pten^+/+^*; *Egfr^+/+^*), *Pten*-MADM (*Pten^-/-^*; *Egfr^+/+^*) and *Pten*-MADM-*Egfr*-cKO (*Pten^-/-^*; *Egfr^-/-^*), and subjected the samples to the protocol for PIP-seq (see Methods for details) (Figure 5A). In a first step for bioinformatics analysis, we confirmed that each genetic paradigm yielded similar numbers of high-quality cells (Figure S8A-S8D). Next, we identified all expected cell-types based on canonical marker gene expression and uniform manifold approximation and projection (UMAP) for dimension reduction (Figure S8E-S8F). For the subsequent analysis, we focused on non-projection neuron cells (Figure S8G) and identified cell-types based on cell type specific marker gene expression (Figure 5B and Figure S8H), as well as cell-cycle scoring for aIPs and RGPs (Figure S8I). We then used Monocle3 to reconstruct trajectories from RGPs to all other postmitotic cell types (Figure 5C). In line with our histological experiments (Figure 4) we found that aIPs, RGPs and astrocytes were overrepresented in both *Pten* and *Pten-Egfr* mutant condition (Figure 5D). We then assessed DEGs in *Pten^-/-^*;*Egfr^-/-^* cells when compared to *Pten^-/-^*;*Egfr^+/+^* by using the MAST method (Finak et al. 2015). We found that RGPs, astrocytes and aIPs present strikingly distinct transcriptomic profiles (Figure 5E), and that a vast majority of DEGs show a high level of cell-type specificity (Figure 5F). Next, we applied the integrative differential expression and gene enrichment analysis [iDEA; (Ma et al. 2020)] pipeline, which revealed highly cell-type specific overrepresentation of key growth factor signaling pathways (Figure 5G), corroborating but also extending our earlier lower resolution bulk sequencing analysis (Figure 3) to the true single cell level.

**Figure 5.**
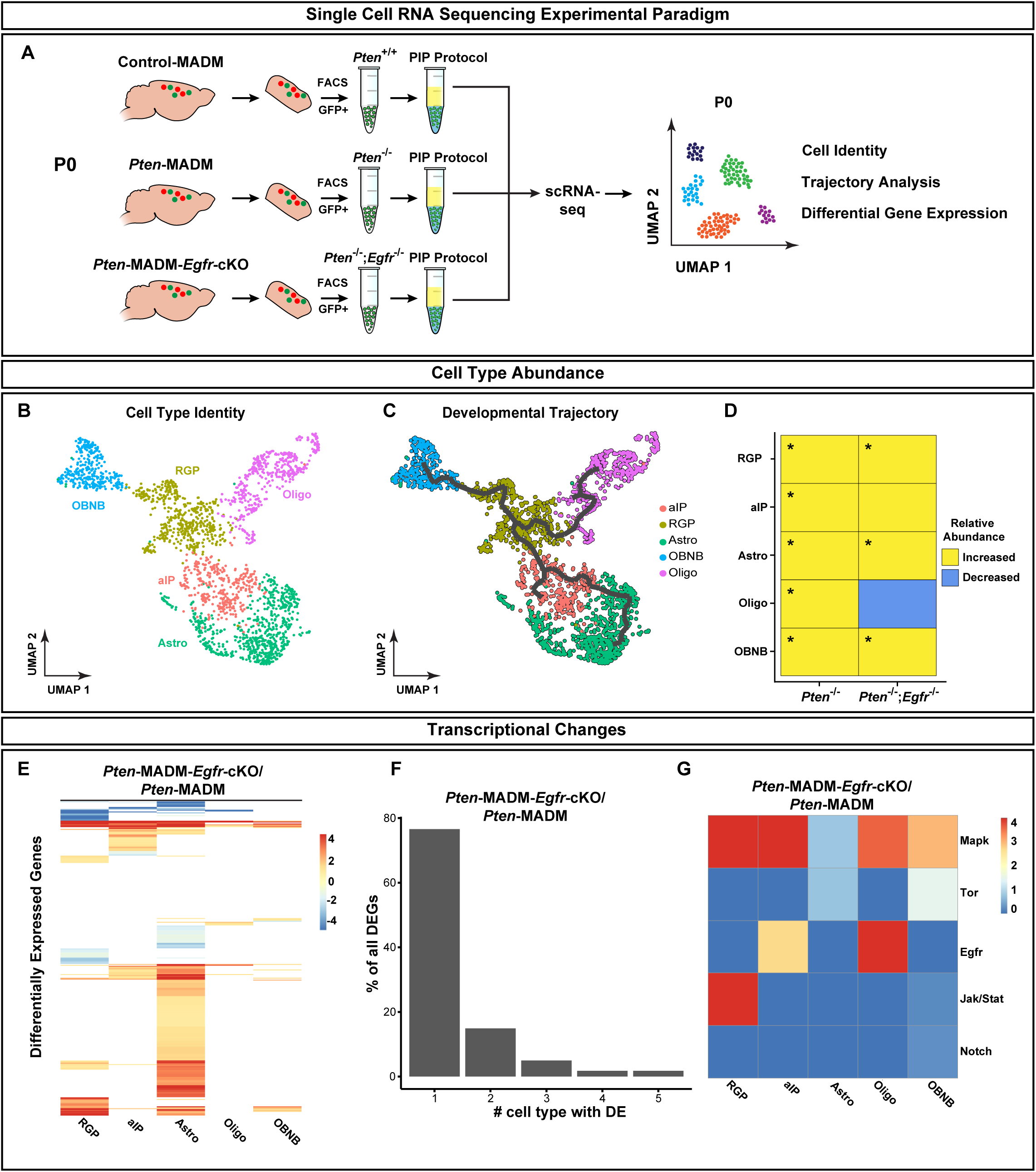
Single cell RNA sequencing reveals highly cell-type specific deregulation of gene expression upon loss of *Pten* and *Egfr*. (A) Schematic overview of the experimental setup for single cell RNA sequencing experiment. Cortex from Control-MADM, *Pten*-MADM and *Pten*-MADM-*Egfr*-cKO was extracted at P0, and GFP^+^ cells were sorted from single cell suspensions, concentrated, and subjected to PIPseq protocol and RNA sequencing. (B) UMAP indicating non-pyramidal neuron cell types from all genotypes. (C) Monocle 3 lineage projection indicating all cell types that originate from the RGP cell population. (D) Cell-type abundance analysis across genotypes. ‘Increased’ refers to an increase in relative abundance for the respective cell type when compared to Control-MADM. ‘Decreased’ refers to a decrease the respective cell type compared to Control-MADM. Chi-square test. * Indicates significance, see Supplemental Table S1 for values. (E) Heatmap indicating DEGs in *Pten*-MADM-*Egfr*-cKO compared to *Pten*-MADM across all cell types. (F) Quantification of % of all DEGs as a function of deregulation in a given number of cell-types. (G) Heatmap indicating integrative differential expression and gene enrichment analysis (iDEA) in *Pten*-MADM-*Egfr*-cKO compared to *Pten*-MADM.

### Epistatic genetic interactions of *Pten* and *Mapk1/3* tune cortical astrocyte production

The combined analysis of bulk and scRNA-seq revealed highly cell-type specific deregulation of genes encoding for major growth factor signaling components, upon loss of *Pten*, besides rendering *Pten^-/-^* mutant cells insensitive to EGFR – one of the most essential growth factor pathways for astrocyte generation. We thus set out to probe further candidates for epistatic interaction with *Pten*, with the goal to get deeper mechanistic insights at the genetic level. We conceived a strategy to combine knockout alleles for selected growth factor signaling components with *Pten*-MADM. The rationale of the assay was to rescue/correct the *Pten* mutant phenotype – and thus to lower overall astrocyte numbers – which would implicate the tested candidate in the causation of the *Pten* phenotype. We first tested genetic interaction separately with *Notch1*, *Stat3*, *Stat5*, *Jak1*, and *Jak2*, which were found high up on the list derived from our RNA sequencing experiments (see above), and which previously have been implicated in astrocyte development. Yet, we could not observe significant rescue of the *Pten* astrocyte overproduction phenotype with any of the above candidates (Figure S9-S12), as was the case with *Egfr*.

Next, we focused on MAPK signaling pathway and conducted a series of epistasis experiments with *Mapk1* (aka *Erk2*) and/or *Mapk3* (aka *Erk1*) in combination with *Pten*-MADM (Figure 6 and S13). First, we generated *Pten*-MADM mosaic in combination with either *Mapk3*-cKO (Figure 6F-G) or *Mapk1*-cKO (Figure 6H-I) but found no significant difference between the elevated level of *Pten^-/-^* mutant astrocytes when compared to *Pten^-/-^*;*Mapk3^-/-^* or *Pten^-/-^*;*Mapk1^-/-^* double mutant condition (Figure 6P). Based on evidence in literature (Li et al. 2012) we next tested for putative functional redundancy between *Mapk3* and *Mapk1*. To this end we generated a series of experimental conditions: *Pten*-MADM with *Mapk3*-cKO and *Mapk1^+/-^*(Figure 6J-6K), *Pten*-MADM with *Mapk1*-cKO and *Mapk3^+/-^* (Figure 6L-6M), and *Pten*-MADM with *Mapk3*-cKO plus *Mapk1*-cKO (Figure 6N-6O), which we compared to *Pten*-MADM and Control-MADM. We found that homozygous elimination of *Mapk3* and having one copy of *Mapk1* in *Pten^-/-^* mutant did not show any difference to the condition with only homozygous *Pten^-/-^* mutant (Figure 6P). In contrast, when we eliminated both copies of *Mapk1* and one or both copies of *Mapk3* together in *Pten^-/-^*, the astrocyte overproduction phenotype was completely rescued and numbers of cortical astrocytes resembled wild-type levels (Figure 6P-6Q). Thus, the elimination of *Pten* results in elevated cortical astrocyte production, independent of *Egfr*, *Notch1*, *Jak1/2*, and *Stat3/3* but via *Mapk1/3*-dependent pathways.

**Figure 6.**
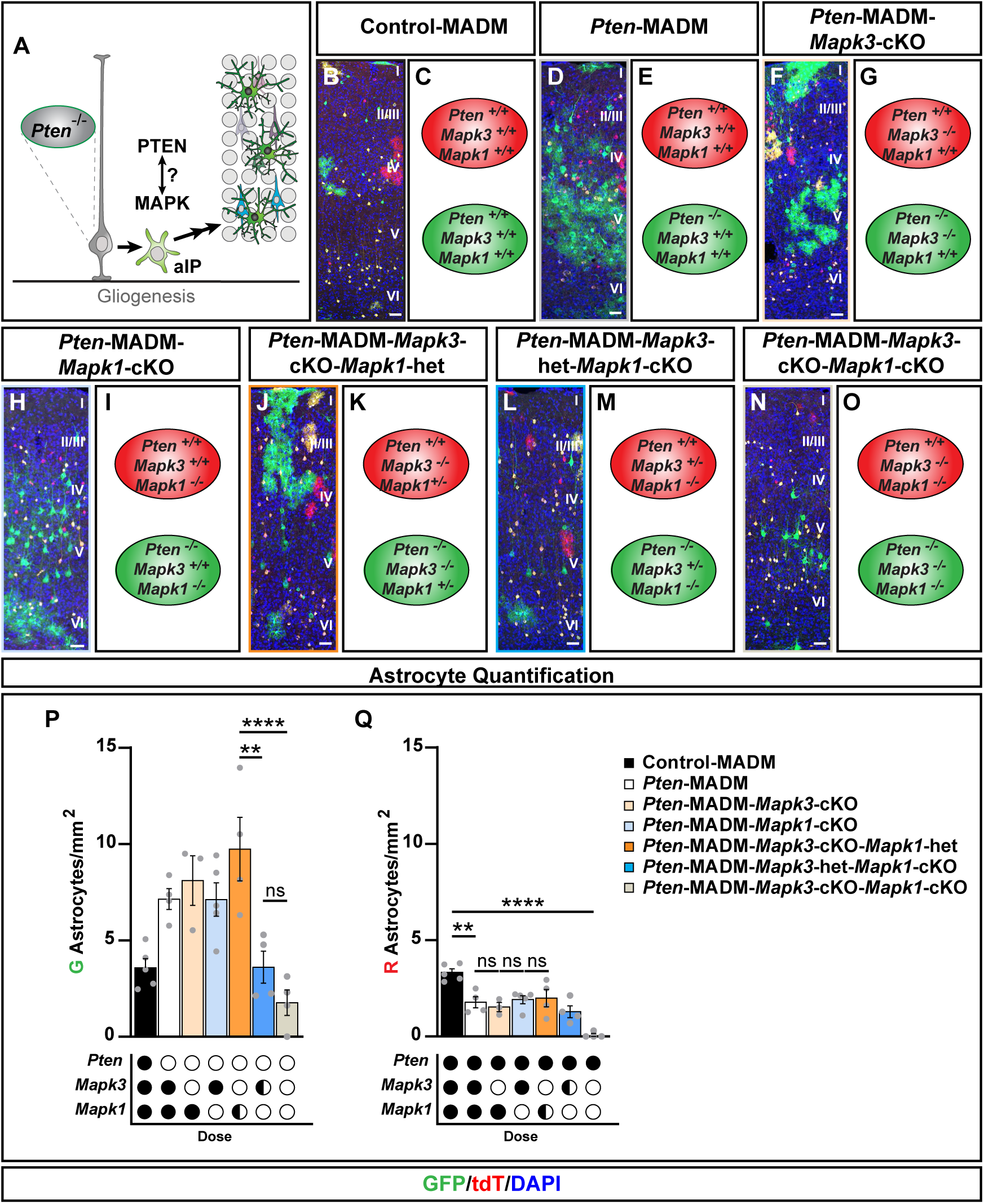
Epistatic genetic interaction of *Pten* and *Mapk1/3*. **(A)** Schematic illustrating putative epistatic interaction between *Pten* and *Mapk* requirement for the production of cortical astrocytes. The concept of the experiment relied on the question whether the *Pten^-/-^*astrocyte overproduction phenotype could be rescued by concomitant deletion of *Mapk*. **(B-O)** Representative images of P21 somatosensory cortex (B, D, F, H, J, L and N), and schematics (C, E, G, I, K, M and O) illustrating the genotype in MADM-labeled GFP^+^ and tdT^+^ cells in Control-MADM (black), *Pten*-MADM (white), *Pten*-MADM-*Mapk3*-cKO (light orange), *Pten*-MADM-*Mapk1*-cKO (light blue), *Pten*-MADM-*Mapk3*-cKO-*Mapk1*-het (orange), *Pten*-MADM-*Mapk3*-het-*Mapk1*-cKO (blue), *Pten*-MADM-*Mapk3*-cKO-*Mapk1*-cKO (beige). (**P and Q**) Quantification of GFP^+^ (P) and tdT^+^ (Q) astrocyte/mm^2^ in the P21 somatosensory cortex in Control-MADM (black), *Pten*-MADM (white), *Pten*-MADM-*Mapk3*-cKO (light orange), *Pten*-MADM-*Mapk1*-cKO (light blue), *Pten*-MADM-*Mapk3*-cKO-*Mapk1*-het (orange), *Pten*-MADM-*Mapk3*-het-*Mapk1*-cKO (blue), *Pten*-MADM-*Mapk3*-cKO-*Mapk1*-cKO (beige). *Pten*, *Mapk3*, and *Mapk1* gene dosage is indicated below each plot. Values represent mean ± SEM. Significance was determined using One-way ANOVA with Tukey’s multiple comparison test. ns = not significant, *p < 0.05, **p < 0.01, ***p < 0.001, ****p < 0.0001. Nuclei were stained using DAPI (blue). Cortical layers are indicated in roman numerals. Scale bar, 50 μm.

## DISCUSSION

The generation of a correctly sized cerebral cortex is a fundamental prerequisite for faithful brain function. Abnormalities in cerebral cortex development that lead to aberrant size may be clinically isolated or could emerge as comorbidities of complex syndromes (Pirozzi et al. 2018; Blumcke et al. 2021; Klingler et al. 2021; Bizzotto and Walsh 2022; Balestrini et al. 2023; Currey et al. 2025). However, while a number of signaling pathways have been linked to regulatory mechanisms controlling brain size, most insight is currently restricted to observations at macroscopic and/or whole tissue level. Here, we focused on the underlying mechanisms associated with macrocephaly and specifically probed the cell-autonomous function of *Pten* in RGP lineage progression with unprecedented single cell resolution (Figure 7). We found that the loss of *Pten* resulted in slightly higher overall projection neuron generation due to increased symmetric progenitor division and enlarged neurogenic potential/neuron output in asymmetrically dividing RGPs. In the absence of *Pten*, aIPs were more numerous and clonal astrocyte output was skewed towards significantly higher clonal units leading to an overabundance of cortical astrocytes. At the mechanistic level, we could show that astrocyte generation was uncoupled from essential EGFR requirement but dependent on intact MAPK signaling. Below we discuss our finding in the context of neurogenic RGP lineage progression and cortical astrocyte ontogenesis.

**Figure 7.**
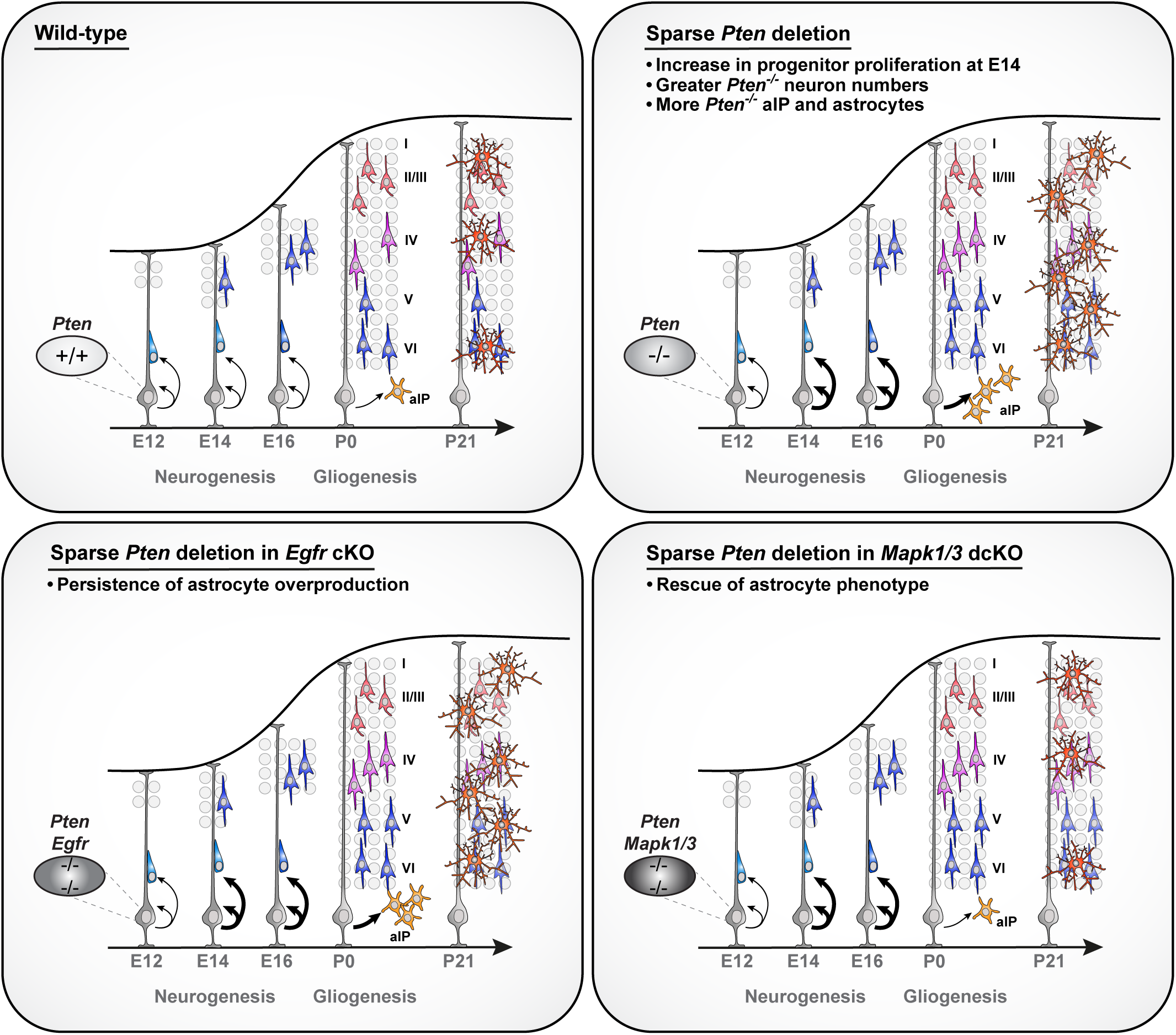
*Pten* modulates RGP lineage progression at distinct stages. Model of RGP lineage progression during cerebral cortex development and the role of *Pten*. (*Top left*) *Wild-type* condition. (*Top right*) Loss of *Pten* amplifies the RGP pool, leading to more progenitors capable of generating a clonal unit. *Pten* mutant progenitors also produces a slightly larger clonal unit than *wild-type* progenitors. In the absence of *Pten*, the number of aIPs was increased and a fraction of RGPs produce abnormally large astrocyte cohorts, leading to an over-abundance of astrocytes throughout the cortex. (*Bottom left*) *Pten* mutant RGPs in combination with *Egfr* loss of function revealed uncoupling of cortical astrocyte production from essential *Egfr* requirement. (*Bottom right*) Epistasis experiments demonstrated that the mutant *Pten* astrocyte overproduction phenotype relied on intact MAPK signaling.

### Role of *Pten* in neurogenic RGP lineage progression

The generation of appropriate numbers of cortical projection neurons by neurogenic RGPs is following a stereotyped quantitative and temporal logic (Lin et al. 2021; Llorca and Marin 2021; Hippenmeyer 2023). The precise switch from symmetric amplifying to asymmetric neurogenic RGP division occurs relatively sharply around E12 in mice (Gao et al. 2014). In our MADM-based clonal analysis we could show that loss of *Pten* not only results in higher single RGP clonal neuron output but also skews the relative proportion of symmetric versus asymmetric division, resulting in slightly but significantly higher progenitor numbers. This *Pten* phenotype did not result in obvious ventricular/subventricular heterotopias as observed in other mutants with increased progenitor numbers, such as for instance in mice carrying mutations in planar cell polarity genes (Hakanen et al. 2019). However, PTEN has in fact been associated in controlling cellular polarity in other contexts but also to some extent in RGPs (Veleva-Rotse and Barnes 2014; Hansen et al. 2017). In effect, the loss of the apical complex protein PALS leads to the absence of nearly the entire cerebral cortex, a phenotype that can be rescued almost completely by concomitant deletion of *Pten* (Lehtinen et al. 2011). Thus, *Pals1* and *Pten* interact genetically to regulate cerebral cortex size. Mechanistically, PALS and PTEN appear to have opposing roles in localizing IGF1R to the apical domain of cortical RGPs, and thereby enabling access to cerebrospinal fluid (CSF)-derived IGFs to modulate progenitor proliferation (Lehtinen et al. 2011). However, the above experiments utilized whole tissue ablation of *Pals*/*Pten*. Thus, global tissue-wide community effects could contribute to the overall RGP phenotype and it will be interesting to assess the precise cell-intrinsic roles of *Pten* and *Pal*s in regulating RGP cell polarity in future experiments. Interestingly, *Pten* has also been shown to genetically interact with *β-catenin*, a key player in the canonical WNT pathway (Chen et al. 2015). These findings are relevant since 1) heterozygous *Pten^+/-^* mice show elevated β-catenin signaling (Chen et al. 2015), 2) mice with elevated β-catenin signaling display brain overgrowth (Chenn and Walsh 2002) and 3) genetic reduction of *β-catenin* may alleviate phenotypes associated with lower *Pten* gene dose (Chen et al. 2015). Collectively, *Pten* appears as a key regulator of RGP polarity, which in turn regulates progenitor proliferation behavior. Yet, it will be important in future studies to not only define the precise biochemical mechanisms, but also to determine the extent of disturbed RGP cell polarity, accompanied by the loss of *Pten*, at the individual progenitor level, and its relevance in cell-autonomously regulating RGP proliferation behavior and neuron output.

The MADM-based clonal analysis has revealed slightly but significantly increased clonal unitary projection neuron output upon loss of *Pten* with a bias towards increased lower layer projection neurons, resonating with a certain level of cell-type specificity of the neuron overproduction phenotype. Intriguingly, deletion of TSC (tuberous sclerosis complex) proteins, also a negative regulator of mTOR signaling and downstream of PTEN/AKT, led to an altered RGP/IP balance with changes of clonal projection neuron unit composition and with increased upper layer neuron generation (Casingal et al. 2025). These findings are in line with recent analysis in the developing superior colliculus where the loss of *Pten* in RGPs resulted in highly cell-type specific over-/underabundance of not only excitatory but also certain types of inhibitory neurons (Cheung et al. 2024). Thus, it is tempting to speculate that distinct components of the PI3K-PTEN-TSC-mTOR axis appear to fulfill highly specific functions during RGP-mediated neuron production, and more generally, that *Pten* function is critical in various RGP stem cell niches in distinct brain regions.

### *Pten* in cortical astrocyte generation

Loss of *Pten* in gliogenic RGPs resulted in increased cortical astrocyte production with enlarged clonal astrocyte units, but how does the absence of *Pten* lead to astrocyte overabundance? While we could not detect any obvious evidence of precocious astrocyte production, the number of aIPs was significantly increased at birth and through the first postnatal week. In any case, previous work has demonstrated that the generation of cortical astrocytes was strictly dependent upon EGFR signaling (Burrows et al. 1997; Sibilia et al. 1998; Beattie et al. 2017; Zhang et al. 2020). Hence, in the absence of EGFR, astrocytes (at least astrocytes derived from *Emx1^+^* RGP lineages) were not present throughout the developing neocortex [this study; (Beattie et al. 2017; Zhang et al. 2020)]. Through genetic epistasis experiments we could show that the loss of *Pten* uncoupled cortical astrocyte production from essential EGFR signaling. These results raised the question: how does the absence of *Pten* trigger exuberant astrocyte production in an EGFR-independent manner. We therefore conducted a series of epistasis experiments with a focus on major signaling hubs that were also identified as significantly deregulated in our scRNA-seq experiments. The rationale behind these experiments was that if a certain deregulated signaling pathway would be overactivated by loss of *Pten* and acting downstream, we should be able to rescue the astrocyte overabundance phenotype. However, when we tested a putative role for *Notch1* or components of the JAK/STAT pathway, all of which have been associated with critical roles in astrocyte generation and/or maturation (Lattke and Guillemot 2022; Markey et al. 2023; Williamson et al. 2025), we could not find a significant lowering of astrocyte generation. In contrast, the concomitant elimination of ERK signaling not only corrected the *Pten* loss of function phenotype but eliminated cortical astrocyte production almost completely, in agreement with previous studies (Li et al. 2012; Gao et al. 2025). We thus conclude that loss of *Pten* renders astrocyte progenitors not only insensitive to EGFR signaling, but also uncouples astrocyte production from NOTCH and JAK/STAT signaling pathways. While *Pten* loss of function condition may reflect a ‘synthetic’ intracellular situation, overriding endogenous, and triggering ectopic signaling pathways, ERK activity and its downstream transcriptional targets likely still operate as a gate keeper for astrocyte generation. It will thus be important in future studies to investigate the specific transcriptional downstream targets translating into quantitatively appropriate astrocyte output. A putative key candidate may be the Pea3 subfamily ETS transcription factor ERM, a well-established FGF downstream target that has been reported to be phosphorylated and transactivated by the MAPK pathway (Tsang and Dawid 2004). In effect, we have also identified *Fgfr* and *Erm*/*Etv5* to be deregulated upon loss of *Pten* in our RNA-seq data sets. However, previous studies have not found macroscopic cortical abnormalities in *Erm* KO mice, albeit astrocyte development has not been assessed (Chen et al. 2005) and should be revisited in detail.

An important aspect about our results concerns the generality of our findings. In our study, we have utilized *Emx1*-Cre driver to target neocortical astrocyte populations. Until recently, it was assumed that all cortical grey matter astrocytes emerge directly or indirectly from *Emx1^+^* RGP lineages. By using TrackerSeq (Bandler et al. 2022) to trace clonally-related astrocytes, a recent study (Zhou et al. 2025) found however that two distinct lineages give rise to cortical astrocytes. Interestingly, only one of the two lineages derived was from *Emx1^+^* RGPs whereas the other did not. It will thus be interesting to assess whether *Pten* function is required in both astrocyte lineages or may be restricted to the *Emx1^+^* lineage in future experiments.

In a broader context, *PTEN* mutations have been implicated in a variety of cancers, some of which arise from glial cell populations (Fenton et al. 2012). Glioblastoma multiforme (GBM) is one of the most common intracranial tumors that arises from astrocytes in adults and is associated with, and often preceded by, *PTEN* LOF (Kwon et al. 2008; Ozawa et al. 2014). Importantly, GBM is frequently aggravated by either EGFR amplification or a hyperactivating mutation (An et al. 2018) but GBM patients are often resistant to receptor-tyrosine kinase inhibitor drugs (RTKI), which inhibit EGFR (An et al. 2018). Our data provides evidence that *Pten* LOF uncouples astrocyte production from EGFR, somehow mirroring the RTKI-resistant GBM condition. Thus, our transcriptome data upon *Pten* and/or *Egfr* deletion could potentially be relevant for future efforts to isolate new pharmalogical targets for patients with RTKI-resistant GBM.

## METHODS

### Husbandry and breeding of mice

All breeding and experiments were performed under a license approved by the Austrian Federal Ministry of Women, Science and Research in accordance with Austrian and EU animal laws (license numbers: BMWFW-66.018/0007-II/3b/2012, BMWFW-66.018/0006-WF/V/3b/2017 and GZ 2025-0.597.515). The mouse colony was maintained following guidelines approved by the institutional ethics committee, the institutional animal care and use committee, and the preclinical facility (PCF) and the Institute of Science and Technology, Austria (ISTA). Mice with specific pathogen free status according to FELASA recommendations (Mahler et al. 2014) were bred and maintained in experimental rodent facilities (room temperature 21 ± 1°C [mean ± SEM]; relative humidity 40%–55%; photoperiod 12L:12D). Food (V1126, Ssniff Spezialitäten GmbH, Soest, Germany) and tap water were available *ad libitum*.

Mouse lines with MADM cassettes inserted in Chr. 19 (Contreras et al. 2021), *Pten*-flox (Groszer et al. 2001), *Egfr*-flox (Lee and Threadgill 2009), *Jak1*-flox (Sakamoto et al. 2016a; Sakamoto et al. 2016b), *Jak2*-flox (Krempler et al. 2004; Wagner et al. 2004), *Stat3*-flox (Moh et al. 2007), *Stat5*-flox (Cui et al. 2004), *Mapk3*-flox (Selcher et al. 2001), *Mapk1*-flox (Samuels et al. 2008), *Notch1*-flox (Yang et al. 2004), *hGFAP-LacZ* (Brenner et al. 1994), *Emx1*-Cre (Gorski et al. 2002), and *Emx1*-CreERT2 (Kessaris et al. 2006) have been described previously.

All histological analyses were carried out in mice in a mixed CD1-C57BL/6J genetic background with the exception of all mice with MADM-19 that had a CD-C57BL/6N background; in both male and female mice. Mice were only selected for desired genotype. No sex specific differences were observed in any experiment. The age and genotype of respective animals is available in the respective figures, legends, and supplementary files.

### Generation of MADM recombinants and experimental mice

In order to generate MADM mosaic mice for the *Pten* gene, the *Pten*-flox allele was recombined into chromosome 19 carrying the *MADM-TG* cassette to generate *MADM-19^TG/TG,Pten^* recombinant stocks. All breeding strategies for the epistasis experiments can be found in the supplemental data figures. In short, the gene of interest (GOI) with flanking *loxP* sites was bred into the existing *MADM-19^TG/TG,Pten^* and *MADM-19^GT/^*^GT^;*Emx1^Cre/+^*stocks (Figures S7, S10-S13).

### Preparation of MADM-labeled tissue

Mouse tissue extraction and preparation were done following previously published protocols (Beattie et al. 2017; Beattie et al. 2020). In short, mice were anesthetized through an intraperitoneal (IP) injection of ketamine/xylazine solution (65 mg, 13 mg/kg body weight, respectively), unresponsiveness was confirmed by pinching the paw. Anesthetized mice were then perfused with ice-cold phosphate buffered saline (PBS) followed by 4% paraformaldehyde (PFA) in PBS. Tissue underwent post-fixation in 4% PFA overnight at 4°C. For embryo tissue collection, pregnant dams were euthanized by cervical dislocation, embryos were collected from uterine horns, and the heads were dissected and fixed in 4% PFA at 4°C for 48 hours. Brains were then washed with PBS and transferred to 30% sucrose for 48-72 hours. Excess sucrose was removed and brains were embedded in Tissue-Tek O.C.T. (Sakura). Embryonic tissue was sectioned at 20μm and directly collected on Superfrost glass slides (Thermo Fisher Scientific). Post-natal tissue with dense MADM labeling was sectioned at 35μm, post-natal tissue with sparse or clonal MADM labeling was sectioned at 45μm. Sections were then collected in 24-well dishes (Greiner Bio-one) and stored at -20°C in a cryoprotectant solution (30% v/v ethylene glycol, 30% v/v glycerol, 10% v/v 0.244M PO4 buffer) until use.

### Immunolabeling of MADM-labeled tissue

Sections on glass slides were first rehydrated 3x with PBS for 15 minutes. Sections were then incubated in blocking solution (10% horse serum, 1% Triton X-100 in PBS) for 1 hr. If necessary, sections underwent antigen retrieval treatment by incubating slides with pre-warmed Citrate Buffer (pH 6) at 85°C for 30 minutes. Primary antibodies were diluted in staining solution (5% horse serum, 0.5% Triton X-100 in PBS) as indicated in the Key Resources Table and 350μl were applied to each slide and incubated overnight at 4°C. The following day, slides were washed 3x with PBS for 5 minutes and incubated with diluted secondary antibodies in staining solution at ambient temperature for 1 hour. Slides were then washed 3x with PBS and incubated with DAPI (1:20,000 in PBS) for 15 minutes. Finally, cover slips were mounted onto the slides with mounting media containing 1,4-diazabicyclooctane (DABCO; Roth) and Mowiol 4-88 (Roth) and stored at 4°C until imaging. Embryonic MADM tissue, up to E18, underwent primary labeling against GFP (Aves Labs) and tdTomato (Sicgen Antibodies). This step is not required for tissue at postnatal timepoints.

BLBP (Chemicon) staining required antigen retrieval using Citrate Buffer (pH 6) at 85°C for 30 minutes. Sections were then incubated for 2 nights with anti-BLBP (rabbit, 1:700 in 5% horse serum, 0.5% Triton X-100 in PBS). Stained sections were washed 2x with PBS for 15 minutes, and one final wash for 1 hour. Sections were incubated with anti-rabbit Alexa Fluor 647 (Molecular Probes) at room temperature for 1 hour. Sections were washed 3x for 5 minutes with PBS, and incubated with DAPI (1:20,000 in PBS) for 15 minutes. Slides were mounted as described above.

### Imaging and analysis of MADM-labeled tissue

Samples were imaged using an LSM800 or LSM900 confocal microscopes (Zeiss). Tiled images of regions of interest were acquired for a minimum of four sections per animal per genotype using either 10x or 20x air objectives. Images were initially processed using Zen Blue software from Zeiss, then imported into Photoshop (Adobe) where the ROI was marked and cell numbers were manually quantified based on marker gene expression. For cell density analysis, ROI was first marked in Photoshop and the area was then measured in the Zeiss Zen Blue software.

### MADM clonal analysis

Clonal analysis using MADM was performed as previously described (Hippenmeyer et al. 2010; Beattie et al. 2020). Briefly, mating cages were set up and females were checked for signs of mating, and subsequently separated from their mates. Pregnant females were injected with 2mg of tamoxifen (TM) (Sigma-Aldrich) dissolved in corn oil (Sigma-Aldrich) at E11, E12, and E13. Embryos were collected by cesarean section of the pregnant dam, followed by hysterectomy, and analyzed at E13 and E16. Cesarean sections, followed by hysterectomy were performed for pregnant dams who did not give birth by E19. Recovered pups were fostered and maintained until analysis at P28. Clone-containing brains were collected, processed and imaged as described above.

### Analysis of neuron and astrocyte morphology

Neuron and astrocyte morphology analysis were done as previously described (Miranda et al. 2024). In short, GFP^+^ cells were imaged using a 63x oil immersion objective with overlapping z-sections. CZI files were converted to IMS using the IMARIS file converter software. IMS files were used in IMARIS to generate 3D reconstructions. The 3D reconstructions were used to measure total cell volume in IMARIS. Sholl analysis was performed on astrocytes to quantify branching complexity.

### Preparation of single-cell suspension and FACS of MADM-labeled tissue

The protocol followed for generating a single-cell suspension and subsequent FACS of MADM-labeled tissue is already published (Laukoter et al. 2020a; Amberg et al. 2024). In short, animals aged P0/P4 were collected and euthanized by decapitation. The cortex was dissected from each brain and placed in cold PBS in 12-well plates while all samples were processed. Single-cell suspensions were generated for individual animals for use as biological replicates. Tissue was dissociated using a kit containing papain with L-cysteine and EDTA (vial 2, Worthington), deoxyribonuclease (DNase) I (vial 3, Worthington), ovomucoid protease inhibitor (vial 4, Worthington), Dulbecco’s modified Eagle’s medium (DMEM)/F12 (Thermo Fisher Scientific), Earle’s Balanced Salts (EBSS) (Thermo Fisher Scientific), horse serum (HS), and fetal bovine serum (FBS). Worthington reagents were reconstituted according to manufacturer’s instructions using the indicated solvents. Isolated cortices were treated with papain-DNase (papain 20 U/ml with 1000 U DNase) and digested for 30 minutes at 37°C in a water bath. Samples then received a solution of EBSS containing 0.67 mg ovomucoid protease inhibitor and DNase (166.7 U/ml) (Solution 2) and the entire solution was triturated to generate a cell suspension and were then centrifuged at 1000 rpm for 5 minutes at ambient temperature. The supernatant was extracted and the remaining cell pellet was resuspended in solution 2. Suspension was then triturated with a p1000 pipette to mechanically dissociate tissue remnants. The suspension was washed with DMEM/F12 and centrifuged at 1500 rpm for 5 minutes at ambient temperature. Cells were resuspended in a final solution of DMEM/F12 with 10 % FBS and 10% HS. Suspensions were kept on ice until sorting.

For GFP^+^ astrocyte sorting, the cell suspensions were incubated for 25 minutes at 37°C in 100µl LacZ staining solution (Abcam, 1:50 dilution). The reaction was terminated by addition of 100 µl of DMEM/F12 with 10% FBS and 10% HS. Cells were kept on ice, shielded from light until sorting was initiated.

Cell sorting using FACS was done as previously described (Laukoter et al. 2020a; Amberg et al. 2024). In short, immediately prior sorting, the cell suspension was strained through a 40µm cell strainer. FACS was performed using a BD FACSAria III, using the 100µm nozzle, cell sample and cell collection devices were kept at a temperature of 4°C. Doublet exclusion was performed to ensure collection of true single cells. For bulk-seq, 500 astrocytes/sample were collected in 4µl cell lysis buffer [0.2% Triton X-100 and RNAse inhibitor (2 U/µl) (CloneTech)]. Samples were transferred into a 96-well plate on dry ice, and stored at -80°C until the plate was full for further processing.

Note: For PIPseq experiments, single-cell suspension was generated in DMEM/F12+RNAse inhibitor (0.2µg/µl) without serum. Cells were sorted into 1.5ml centrifuge tubes with 1ml cell suspension buffer (PIPseq kit FBS-SCR-T2-8-V4.05-3) (Fluent Biosciences). To concentrate cell suspension, centrifuge tubes were spun down for 10 minutes at 400 rpm, supernatant was removed until approximately 10µl remained. Cells were resuspended, and 5µl of the cell suspension were counted on a Nucleocounter NC250 (Chemometec). The appropriate volume of the remaining 5µl, depending on cell concentration, was used as input for the PIPseq protocol (target of 1000 cells/ µl, max input per PIP tube is 4000 cells in 4µl). The PIP protocol was followed according to the kit instructions without further modification.

### Processing and statistical analysis of bulk RNA-seq data on enriched astrocyte samples

Bulk RNA-Seq was performed by sorting 30-400 cells in SMART-Seq3 buffer (Hagemann-Jensen et al. 2020). Library production and sequencing (Illumina NovaSeqSP platform) was performed at VBCF GmbH (Vienna, https://www.viennabiocenter.org/vbcf/).

Reads were aligned to GRCm39 with Gencode M27 annotation (downloaded from https://www.gencodegenes.org) using STAR aligner. We removed 3 samples due to low alignment rate <45% and 2 samples due to low correlation with biological replicates. Finally, we performed principal component analysis (PCA) of the top 500 variable genes in all analyzed samples and removed 3 samples based on its position in the PCA plot. PCA plot of final samples is shown in Supplemental Data Figure S6A. We retained 3-5 replicates per sample, which showed a unique alignment rate between 46% - 70% resulting in 1.9M–3.5M uniquely aligned reads for downstream analyses.

The conditional deletion allele removes *Pten* exon 5 (Groszer et al. 2001). Therefore, we quantified reads in this region (chr19:32777198-32777560, GRCm39) to test for efficient recombination as shown in Supplemental Data Figure S6B. Differential gene expression was calculated using DESeq2 separately for each developmental time point using contrasts to compare pairwise comparisons of genotypes. Only genes with a mean read count >10 were used for further analysis. Figure 3B, C: Number and overlap statistics of differentially expressed genes (DEG) from comparison of *Pten*-MADM/Control-MADM with an adjusted p-value <0.01 and a log2 fold-change >0 (up, higher in respective *Pten* deletion cells compared to Control-MADM cells) or <0 (down, lower in respective *Pten* deletion cells compared to Control-MADM cells). Gene Ontology (GO) enrichment analysis was performed on DEGs (adjusted p-value < 0.01) using enrichGO function from the clusterProfiler package. Figure 3D: DEGs connected to the GO term gliogenesis (GO:0042063) were extracted and log2 fold change (as determined by DESeq2) displayed.

### STRING network analysis

For STRING network analysis, all significant DEGs (p-value < 0.05) were uploaded, along with the genes listed in Figure 3E, to the STRING website (https://string-db.org/) via the ‘Multiple protein’ function and was selected from the ‘Organisms’ drop-down menu. In the settings tab of the network, disconnected nodes were excluded by selecting ‘hid disconnected nodes in the network’; all other settings were left at their default parameters. Networks were then exported and processed for figures using Cytoscape.

### Library preparation for single cell RNA sequencing

Library preparation was performed using Fluent Bioscience’s PIPseq T2 Single Cell RNA kit v4.0PLUS (FB0001026) according to the manufacturer’s protocol. Libraries were pooled and sequenced on an Illumina NovaSeqX platform at the VBCF NGS Unit (https://www.viennabiocenter.org/facilities/). Initial alignment and preparation of count matrices was performed using pipseeker-v3.3.0 and pipseeker-gex-reference-GRCm39-2022.04. Downstream analyses were performed using R v4.3.2 and Seurat v5.0.1.

### Single cell RNA sequencing analysis

Supplemental Data Figure S8A-S8D: Data was filtered (parameters: nFeature_RNA > 500 & nFeature_RNA < 6000 & nCount_RNA < 40000 & percent.mt < 15), and processed using standard pipelines. Supplemental Data Figure S8E: UMAP was performed using RunUMAP with dims = 1:25, clusters were determined using FindNeighbors with dims = 1:25 and FindClusters with resolution = 0.3. This analysis identified 7 clusters. Cells from one cluster were not well localized on the UMAP, did not show specific marker genes and were removed (39 cells, 0.6% of total cells). Supplemental Data Figure S8I: Cell cycle scoring was performed using Seurat’s CellCycleScoring function with genes from https://github.com/hbc/tinyatlas/tree/master/cell_cycle. The difference between G2M and S phase scores was determined and regressed out during scaling.

Figure 5C: Trajectory analysis was performed using monocle3 (v1.3.7). Figure 5D: Significance of changes in relative abundance was determined using chi squared test. Differential expression between cell types was calculated using MAST (v1.28.0) (Finak et al. 2015). To draw DEG heatmaps adjusted p-values of significant DEGs (adjusted p < 0.1) were transformed to a score (log10 (adjusted p) *-1) and visualized using pheatmap (v1.0.12). Gene Ontology (GO) term analysis was performed using iDEA (v1.0.1) (Ma et al. 2020) on selected GO terms and differential expression statistics (as above). The p-values, as determined by iDEA, were corrected for multiple testing using p.adjust with standard parameters. GO terms were filtered for significant enrichment and grouped using simplifyEnrichment (v1.12.0) (Gu and Hubschmann 2023) in a 2-step process. We used the uncorrected p-values to allow for a comprehensive documentation of changes. In the first step, broad GO groups were determined, including a group containing major signaling pathways. GO terms in the signaling pathway group were grouped in a second step using simplifyEnrichment. The smallest adjusted p-value was used for plotting the heatmap.

### Quantification and statistical analysis

Statistical analyses were performed in the Graphpad Prism v10 software. Raw data expressed as proportions was transformed using the Y=arcsin(√Y) formula in order to stabilize variance and prepare for statistical testing. Statistical tests used include: variants of two-tailed unpaired t-test for comparisons between two groups, one-way ANOVA for comparisons between multiple groups or two-way ANOVA for comparisons between groups split by two variables (i.e. time and genotype). For G/R quantifications: A biological sample was defined as the G/R ratio from one animal resulting from the quantification of at least 5 sections/animal in adults and 5-10 sections/animal in embryonic stages. For cell density (cells/mm^2^) quantifications: A biological sample was defined as the cell density from an animal resulting from the quantification of at least 5 sections/animal in adults [i.e. the number of all GFP^+^ cells in animal “A”/ the sum of the area from all quantified regions in animal “A”]. All figures include information about the test used, as well as the corresponding p-value and other relevant data.

## ACKNOWLEDGMENTS

We thank Kay-Uwe Wagner (Wayne State University) for generously sharing *Jak1/2*–flox mouse lines; A. Sommer (VBCF GmbH, NGS Unit) for technical support; N. Kim, V. Mick, S. Schnabl, S. Gobeil, and L. Andersen for technical assistance; all members of the Hippenmeyer lab for discussion and B. Novitch for comments on earlier versions of the manuscript. This research was supported by the Scientific Service Units (SSU) of IST Austria through resources provided by the Imaging and Optics Facility (IOF), Lab Support-(LSF) and Preclinical Facilities (PCF). O.A.M received support from the Austrian Academy of Sciences ÖAW (DOC 186584), and N.A. from FWF Elise Richter Program (Grant V1041T). This work was also supported by IST Austria institutional funds; FWF SFB F78 (Neuro Stem Modulation) to S.H., and the European Research Council (ERC) under the European Union’s Horizon 2020 research and innovation programme (grant agreement No 725780 LinPro) to S.H.

## Author Contributions

S.H. conceived the research. S.H., O.A.M., X.C. and F.M.P. designed all experiments and interpreted the data. O.A.M., X.C., F.M.P., A.D., N.A., A.V., A.H. and C.S. performed all the experiments. F.M.P. and O.A.M. performed all computational and bioinformatics analysis with inputs from A.V. and S.H. C.M. and B.H provided critical mutant mouse strains, and T.R. was instrumental in the generation of MADM mouse models. S.H. and O.A.M. wrote the manuscript with inputs from A.V. and F.M.P. All authors edited and proofread the manuscript.

## Materials Availability

All published reagents and mouse lines will be shared upon request within the limits of the respective material transfer agreements.

## Data and Code Availability

All data generated and analyzed in this study are included in the paper and/or supplementary materials. Raw sequencing data will be submitted to Gene Expression Omnibus (GEO).

## SUPPLEMENTARY FIGURE LEGENDS

**Figure S1.**
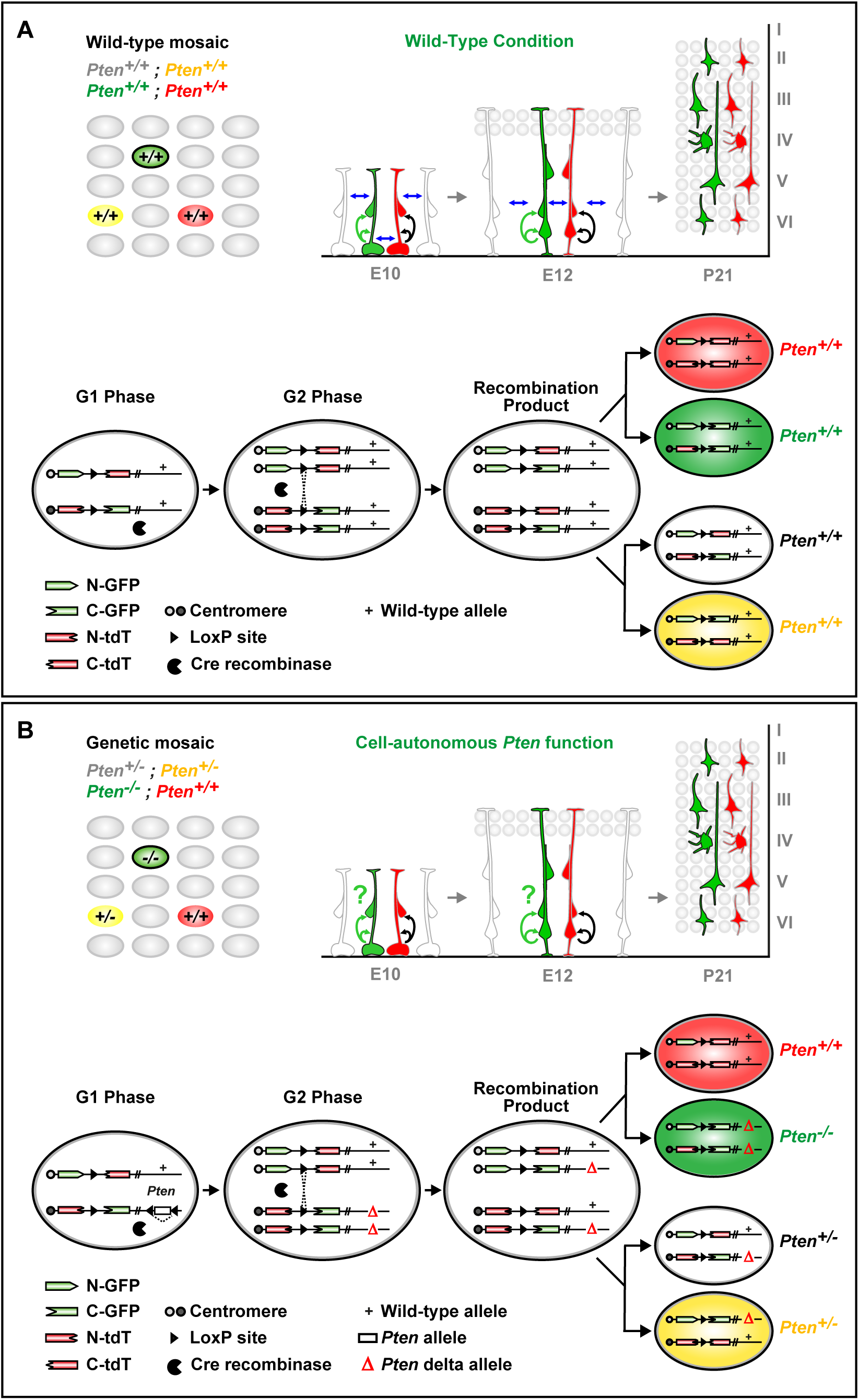
Genetic Paradigms for Wild-Type and *Pten* Sparse Deletion using MADM. **(A)** (*Top*) Illustration of sparse MADM labeling in wild-type mosaic MADM animals. All cells, regardless of fluorescent labeling show identical genotypes. (*Bottom*) Schematic representation of MADM principle to generate sparse labeling. MADM relies on Cre-mediated interchromosomal recombination of the two split marker genes on homologous chromosomes. **(B)** (*Top*) Illustration of sparse *Pten* deletion using MADM, generating two distinct genetic lineages labeled in GFP (*Pten*^-/-^) and tdTomato (*Pten*^+/+^), all other cells, including unlabeled and yellow cells, are heterozygous for *Pten*. Sparse ablation of *Pten* bypasses lethality observed in *Pten* KO and cKO, thereby enabling longitudinal studies not possible with other conditional knockout animal models. (*Bottom*) Schematic representation of MADM principle to generate a sparse genetic *Pten* mosaic. The *Pten-flox* allele was introduced distal to the *TG* MADM cassette via meiotic recombination. Cre recombinase mediates *cis*-recombination of the *loxP* sites of the conditional *Pten* allele, removing exon 5 resulting in a *Pten*-Δ allele, as well as *trans*-recombination between the MADM cassettes. The resulting daughter cells can be either a red *Pten*^+/+^ cell and a green *Pten*^-/-^ cell (upper branch) or a yellow *Pten*^+/-^ cell and an unlabeled *Pten*^+/-^ cell (lower branch). Schematic is reused, adapted and modified from (Beattie et al. 2017), with permission.

**Figure S2.**
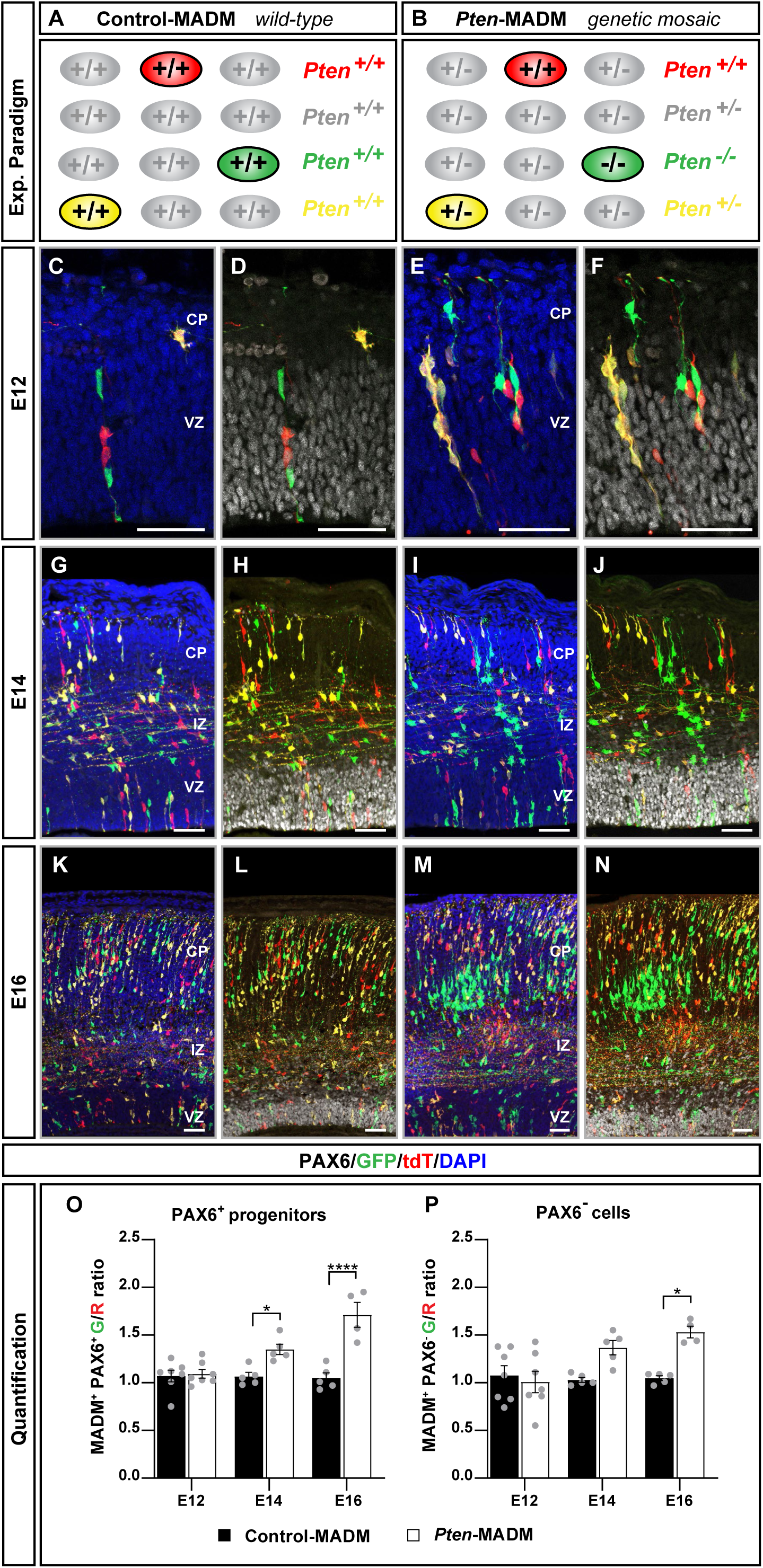
*Pten* deletion leads to expansion of the cortical RGP pool during embryogenesis. **(A-B)** Schematic depicting cellular genotypes in Control-MADM (A, all cells have intact *Pten* alleles) and *Pten*-MADM (B, red tdT^+^ cells have intact *Pten* alleles, green GFP^+^ cells lack functional *Pten*, and yellow tdT^+^/GFP^+^ cells are heterozygous for *Pten*. **(C-N)** Representative confocal images depicting PAX6^+^ and MADM-labeled cells in the developing cortical plate in Control-MADM (C, D, G, H, K and L) and *Pten*-MADM (E, F, I, J, M and N) at E12 (C-F), E14 (G-J), and E16 (K-N). (**O-P**) Quantification of PAX6^+^ progenitors (O) and PAX6^-^ cells (P) at E12, E14, and E16 in Control-MADM (black bars) and *Pten*-MADM (white bars). Values represent mean ± SEM. Significance was determined using two-way ANOVA with Šídák’s multiple comparison test. *p < 0.05, ****p < 0.0001. Nuclei were stained using DAPI (blue). VZ, ventricular zone; IZ, intermediate zone; CP, cortical plate. Scale bar, 50 µm.

**Figure S3.**
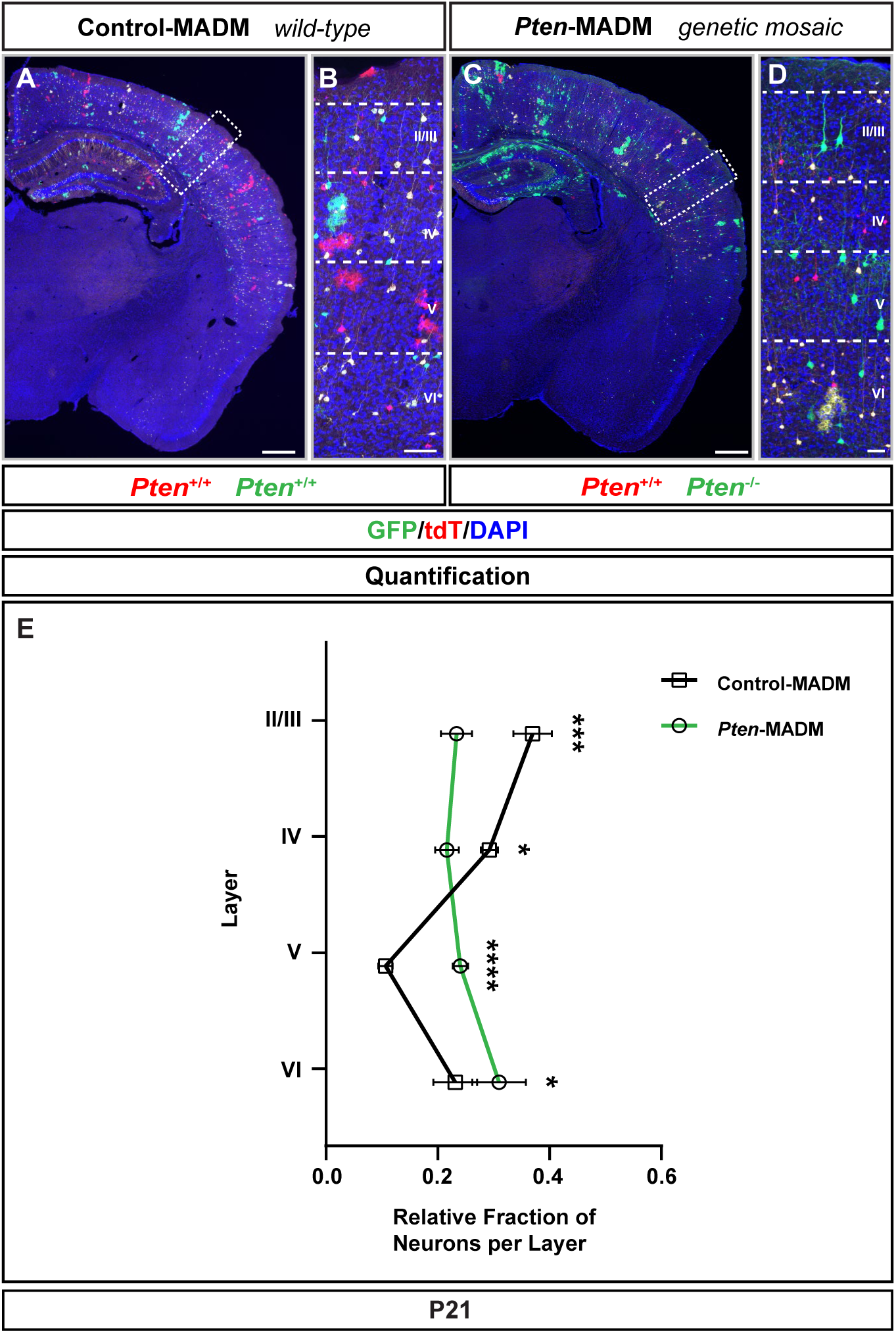
*Pten* mutant neurons differ in abundance across the cortical laminae. **(A-D)** Representative images of somatosensory cortex in overview (A and C) and higher resolution of cortical plate (B and D) in Control-MADM with red tdT^+^ *Pten^+/+^* cells and green GFP^+^ *Pten^+/+^* cells, and *Pten*-MADM with red tdT^+^ *Pten^+/+^* cells and green GFP^+^ *Pten^-/-^* cells at P21. (**E**) Quantification of the relative fraction of GFP^+^ neurons in Control-MADM (black line) and *Pten*-MADM (green line) across the cortical laminae at P21. Nuclei were stained using DAPI (blue). Cortical layers are indicated (dotted line) in roman numerals. Values represent mean ± SEM. Significance was determined using two-way ANOVA with Šídák’s multiple comparison test. *p < 0.05, ***p < 0.001, ****p < 0.0001. Scale bar, 500 μm (A and C), 50 μm (B and D).

**Figure S4.**
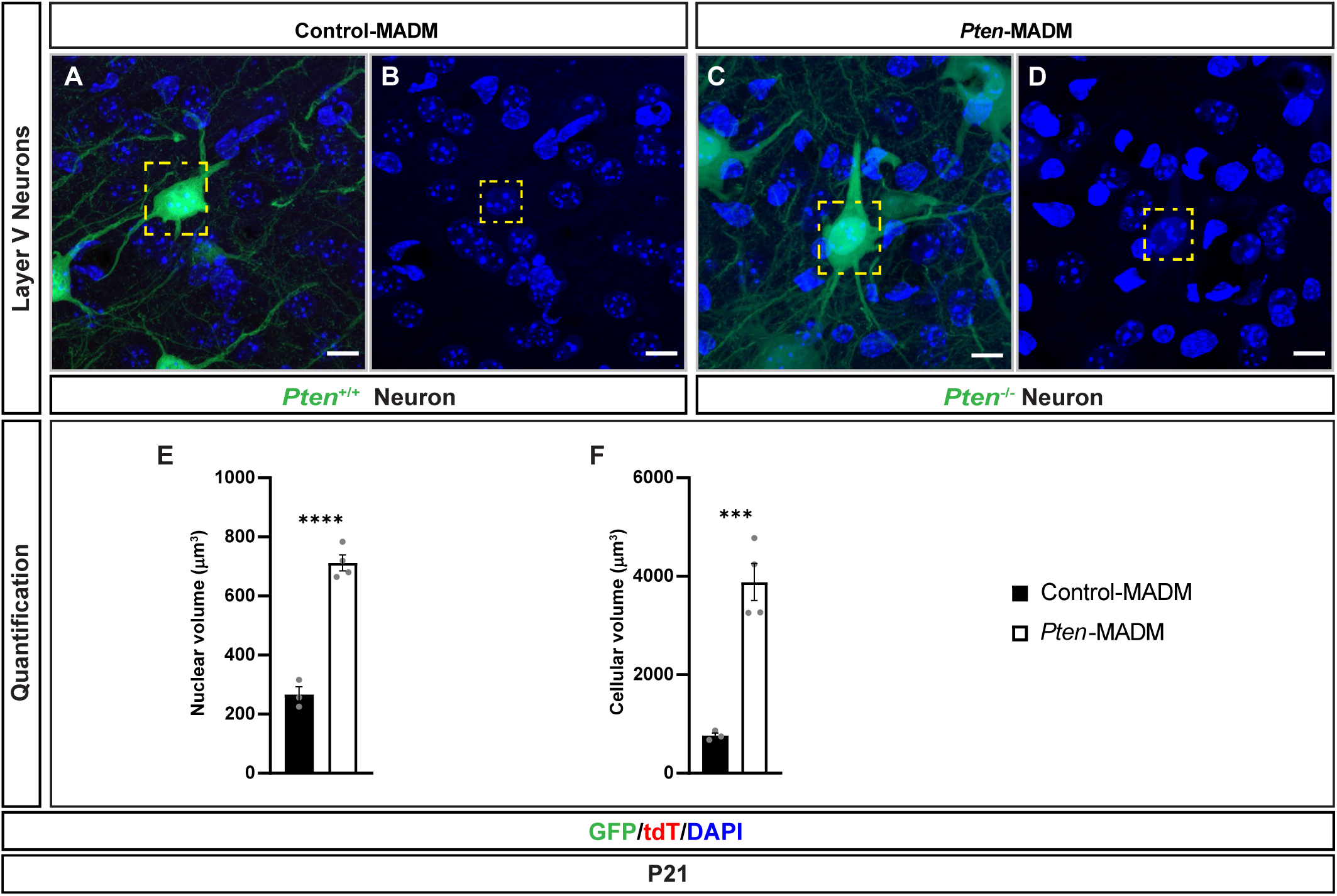
*Pten* cell-autonomously controls neuron cell size. **(A-D)** Representative images of GFP^+^ layer V projection neurons in Control-MADM (A and B, *Pten^+/+^*) and *Pten*-MADM (C and D, *Pten^-/-^*) cells at P21. Cell bodies (A, C) and nuclei (B, D) are indicated by yellow dotted box. (**E-F**) Quantification of cellular soma (E) and nuclear (F) volumes in GFP^+^ neurons in Control-MADM (black bars) and in *Pten*-MADM (white bars). Nuclei were stained using DAPI (blue). Values represent mean ± SEM. Significance was determined using two-tailed unpaired t-test. ***p<0.001, ****p<0.0001. Scale bar, 5 μm.

**Figure S5.**
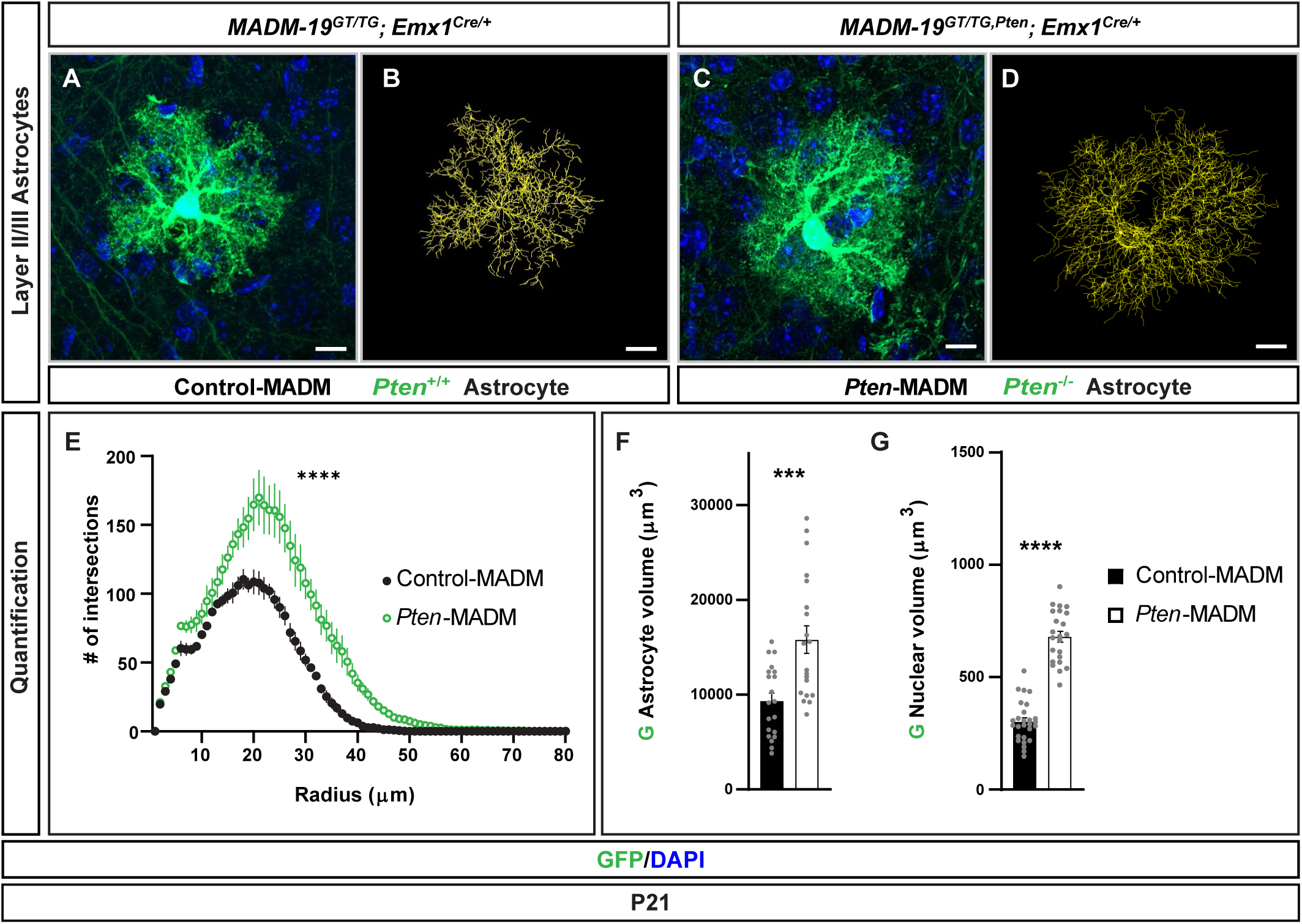
Astrocyte morphology is significantly altered by *Pten* deletion. **(A-D)** Representative images of GFP^+^ layer II/III cortical astrocytes (A and C) and Imaris-based astrocyte processes tracing reconstructions (B and D) in Control-MADM (A and B, *Pten^+/+^*) and *Pten*-MADM (C and D, *Pten^-/-^*) cells at P21. (**E**) Sholl analysis of layer II/III cortical astrocytes in Control-MADM (black; *Pten^+/+^*) and *Pten*-MADM (white, *Pten^-/-^*) cells at P21. (**F-G**) Quantification of total astrocyte (F) and nuclear (G) volumes in GFP^+^ astrocytes in Control-MADM (black bars) and in *Pten*-MADM (white bars). Nuclei were stained using DAPI (blue). Values represent mean ± SEM. Significance was determined using two-way ANOVA (E) and two-tailed unpaired t-test (F and G). ***p<0.001, ****p<0.0001.

**Figure S6.**
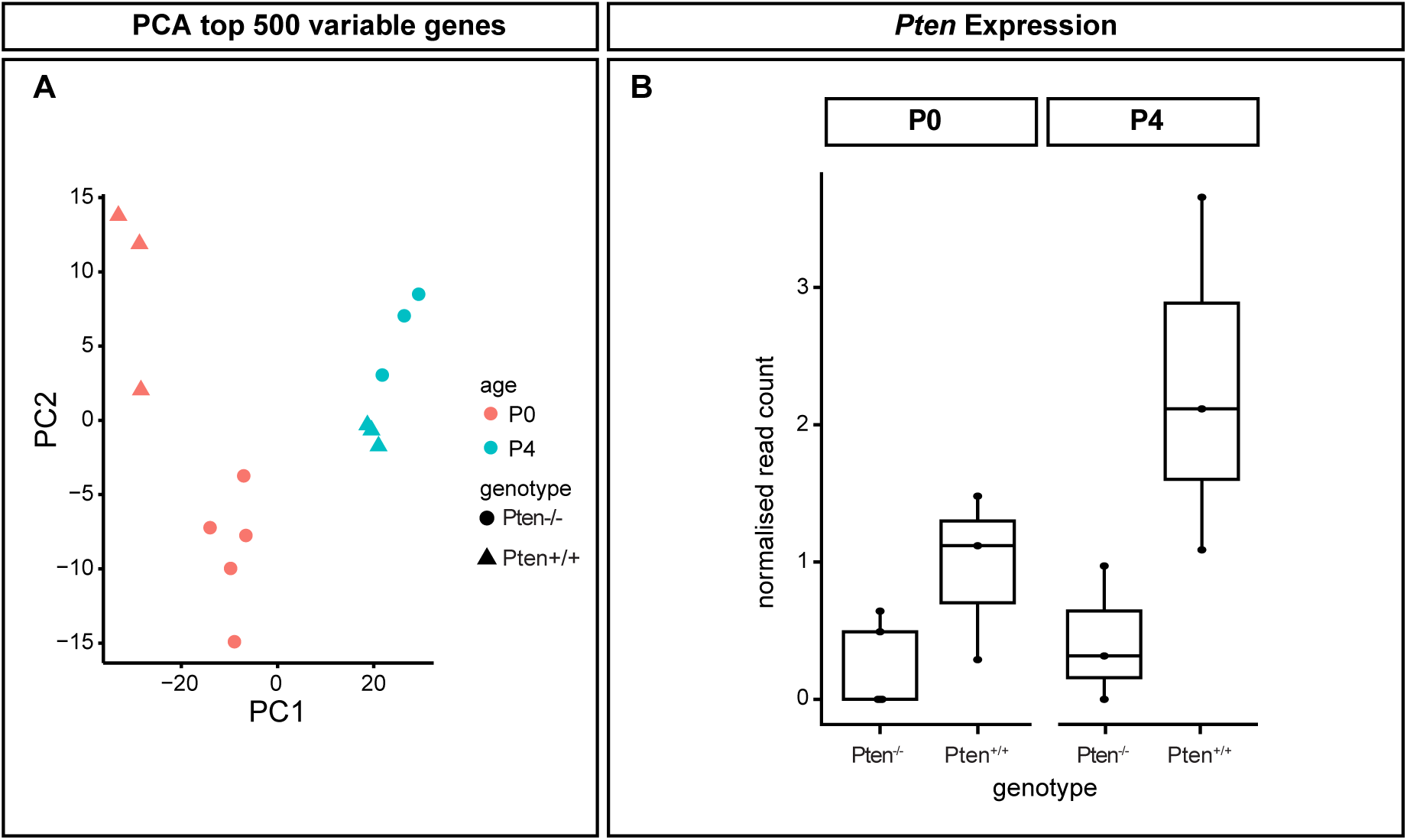
Bulk-RNA-seq of highly pure MADM-labeled astrocytes. (A) Principle Component Analysis (PCA) plot for all samples at P0 and P4. P0 samples segregate into two clusters based on genotype, while P4 cluster closer together, indicating a diminishing difference in transcriptome. (B) Transcript levels for *Pten* Exon 5 in *Pten*^+/+^ and *Pten*^-/-^ astrocytes at P0 and P4.

**Figure S7.**
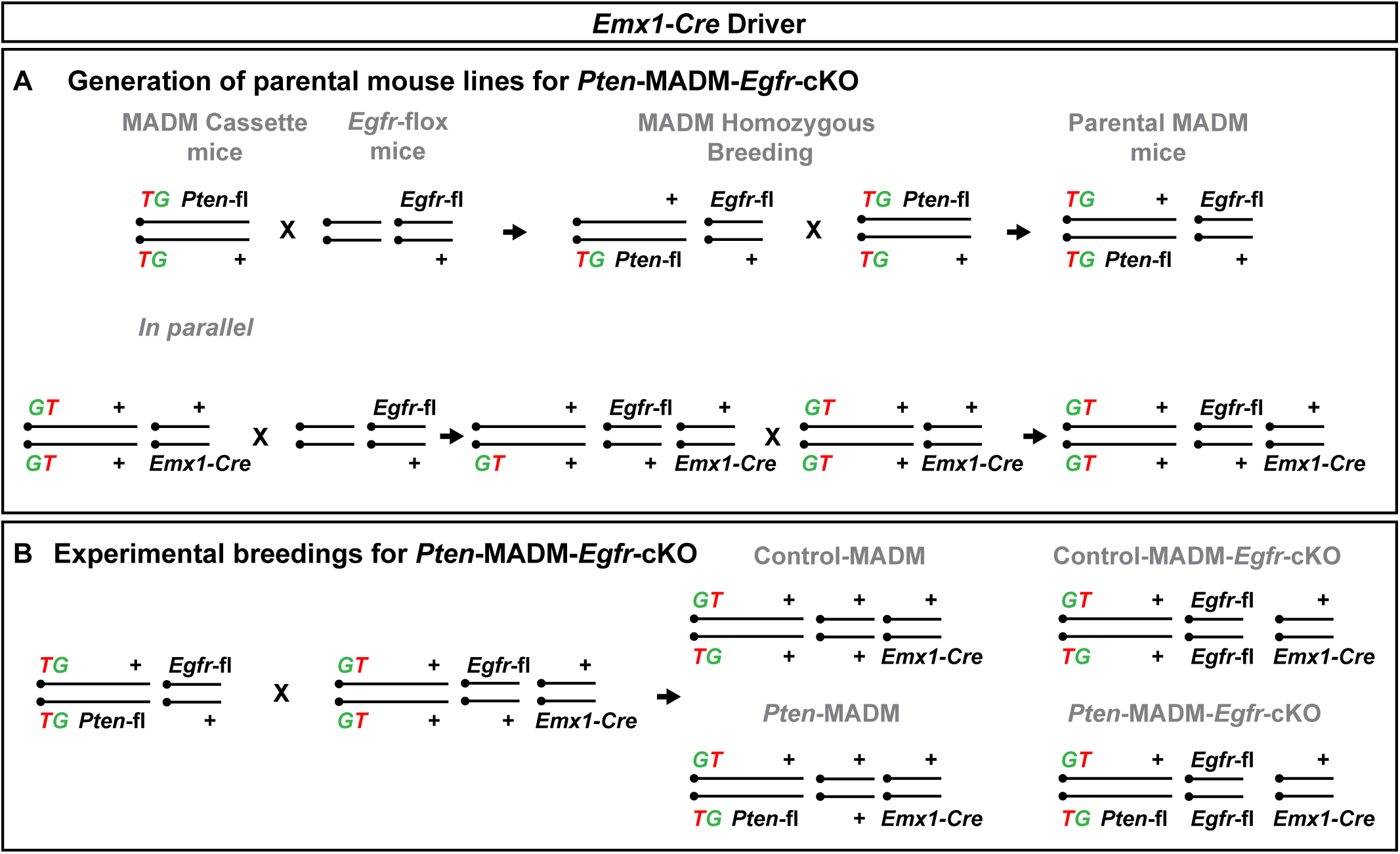
Breeding scheme for generation of mosaic *Pten*-MADM in *Egfr*-cKO background. (A) Parental breeding scheme. *Egfr*-flox mice were crossed to *MADM-19^TG/TG,Pten^* and *MADM-19^GT/GT^*;*Emx1^Cre/+^*. The F_1_ offspring were heterozygous for the respective MADM cassette (*TG-Pten* or *GT*), *Emx1*-Cre, and *Egfr*-flox. These F_1_ offspring were crossed with *MADM-19^TG/TG,Pten^* and *MADM-19^GT/GT^*;*Emx1^Cre/+^*, respectively, to obtain MADM cassette homozygosity. The F_2_ offspring were used for strain maintenance and experimental breeding. Note: It is possible to carry out a further crossing between F_2_ and *MADM-19^GT/GT^*;*Emx1^Cre/+^* to achieve Cre homozygosity. (B) F_2_ offspring carrying MADM cassettes and *Egfr-*flox alleles were crossed to generate all four experimental genotypes as indicated.

**Figure S8.**
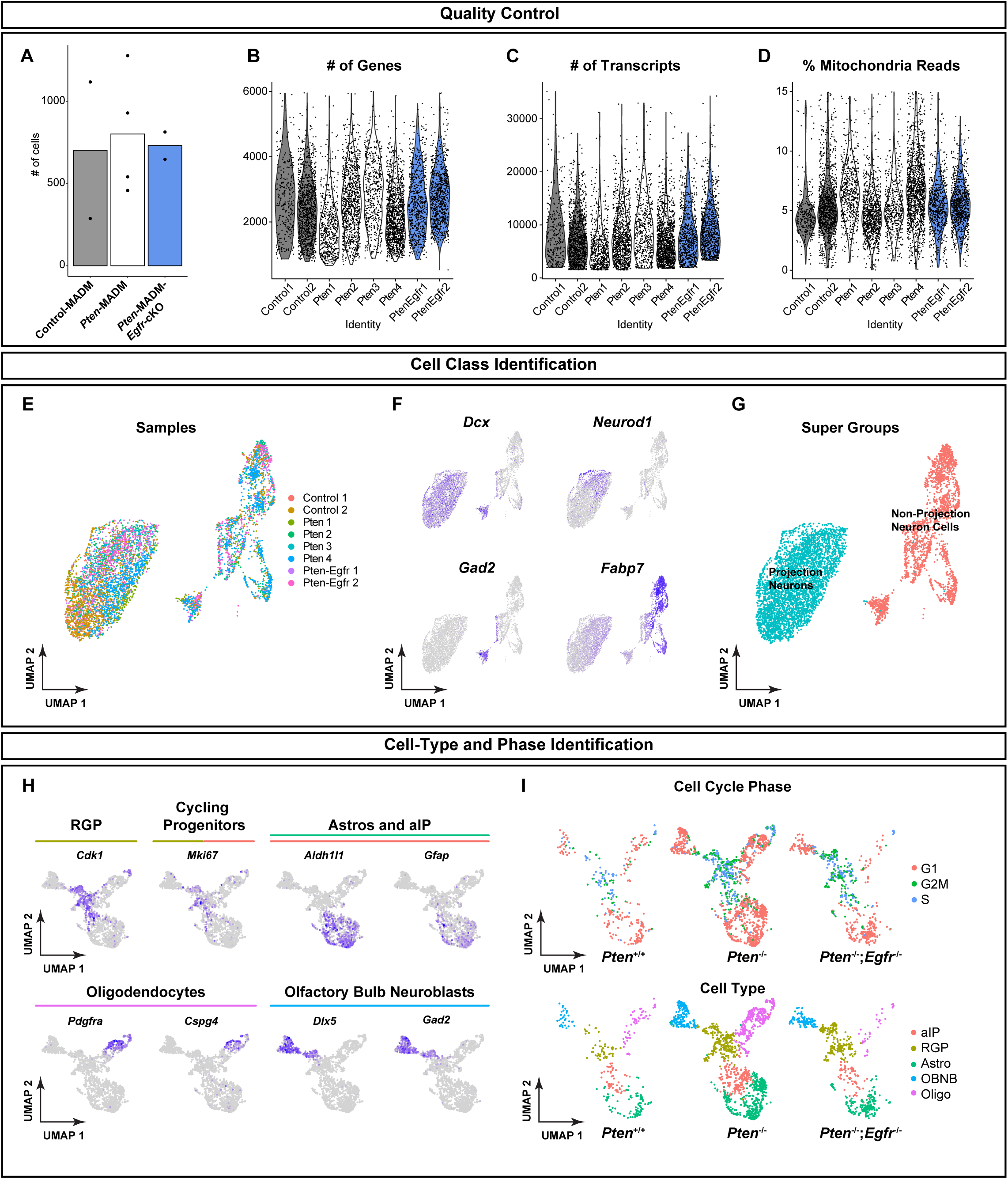
scRNA-seq analysis of P0 cortex from Control-MADM, *Pten*-MADM, and *Pten*-MADM-*Egfr*-cKO animals. (A) Bar plot showing similar cell number averages, collected across the three genotypes, as well as individual samples as data points on each graph. (B) Number of unique gene reads per cell in each sample. (C) Counts of individual transcripts detected per cell across samples. (D) Percent of mitochondrial RNA detected per cell across all samples. (E) UMAP of all cells from all genotypes. (F) UMAPs for identifying broad cell classes by expression of *Dcx* for neurons, *Neurod1* for immature neurons, *Gad2* for olfactory bulb neuroblasts, *Fabp7* for glial cell types. (G) Classification of projection neurons and non-projection neuron cells based on analysis in S8F. (H) Expression of cell-type markers for RGPs (*Cdk1*), cycling progenitors (*Mki67*), astrocytes and progenitors (*Aldh1l1* and *Gfap*), oligodendrocytes (*Pdgfra* and *Cspg4*), and olfactory bulb neuroblasts (*Gad2* and *Dlx5*). (I) UMAPs of cell cycle phase of cells in each genotype (top) and final cell type annotation for each genotype (bottom).

**Figure S9.**
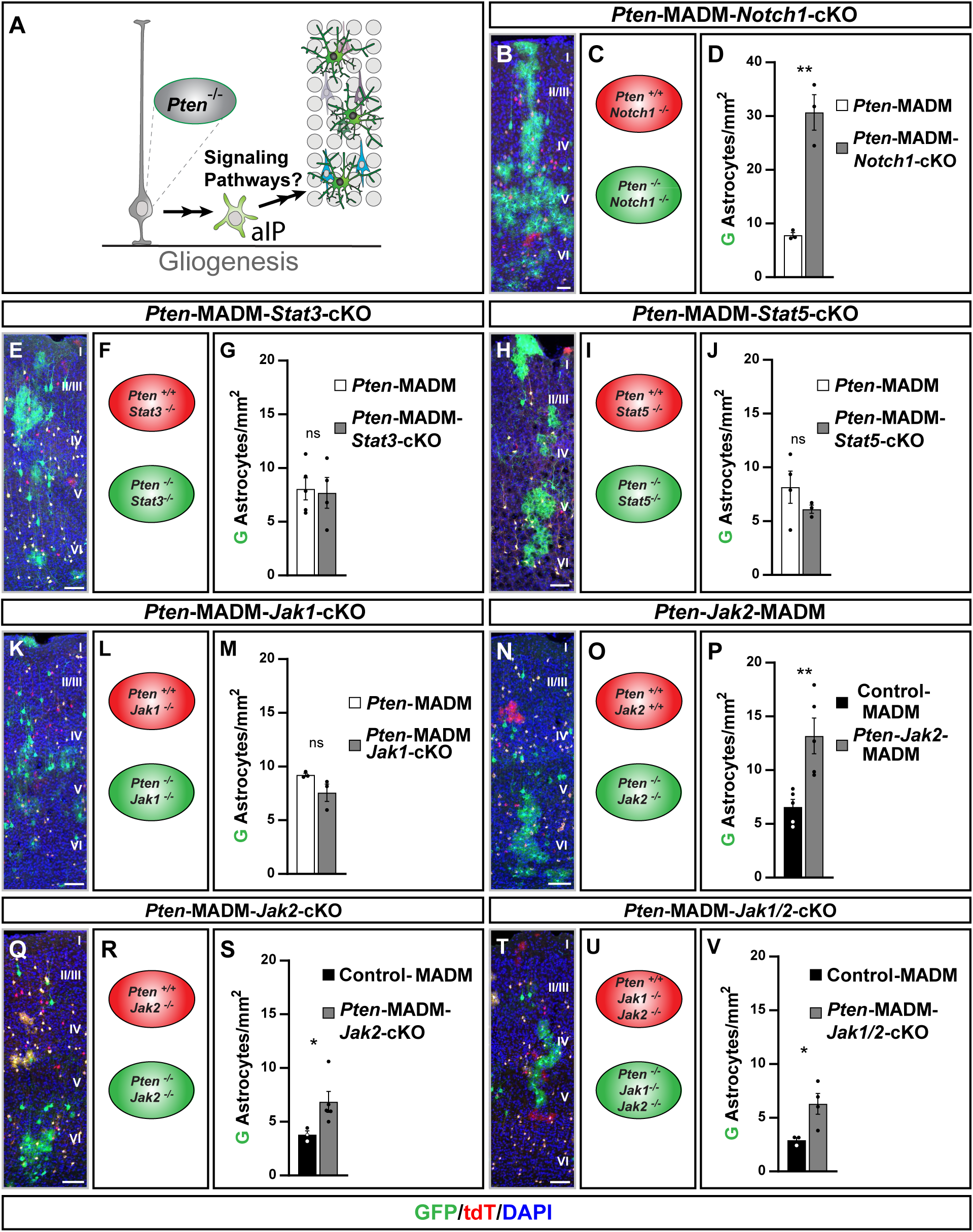
*Pten^-/-^* mutant astrocytes expand independently of *Jak*/*Stat* and *Notch* signaling. (A) Schematic illustrating putative epistatic interaction(s) between *Pten* and distinct major signaling pathways involved in the production of cortical astrocytes. The concept of the experiment relied on the question whether the *Pten^-/-^* astrocyte overproduction phenotype could be rescued by concomitant deletion of *Notch1*, *Stat3/5* or *Jak1/2*. **(B-V)** Representative images of P21 somatosensory cortex (B, E, H, K, N, Q and T), schematics illustrating the respective genotype in MADM-labeled GFP^+^ and tdT^+^ cells (C, F, I, L, O, R and U) and quantification of GFP^+^ astrocyte/mm^2^ (D, G, J, M, P, S and V) in *Pten*-MADM-*Notch1*-cKO (B-D), *Pten*-MADM-*Stat3*-cKO (E-G), *Pten*-MADM-*Stat5*-cKO (H-J), *Pten*-MADM-*Jak1*-cKO (K-M), *Pten*-*Jak2*-MADM (N-P), *Pten*-MADM-*Jak2*-cKO (Q-S), and *Pten*-MADM-*Jak1/2*-cKO (T-V). Note that astrocyte/mm^2^ was significantly higher than in Control-MADM (P, S and V), or not significantly different when compared to *Pten*-MADM (G, J and M; i.e. elevated when compared to Control-MADM), except in *Pten*-MADM-*Notch1*-cKO where the number of astrocyte/mm^2^ was even higher than in *Pten*-MADM (D). Nuclei were stained using DAPI (blue). Cortical layers are indicated in roman numerals. Values represent mean ± SEM. Significance was determined using unpaired t-test (D, J, M, P), Mann-Whitney test (G), Welch’s t-test (S, V). ns not significant, *p<0.05, **p<0.01. Scale bar, 50 μm (B, E, H, K, N, Q and T).

**Figure S10.**
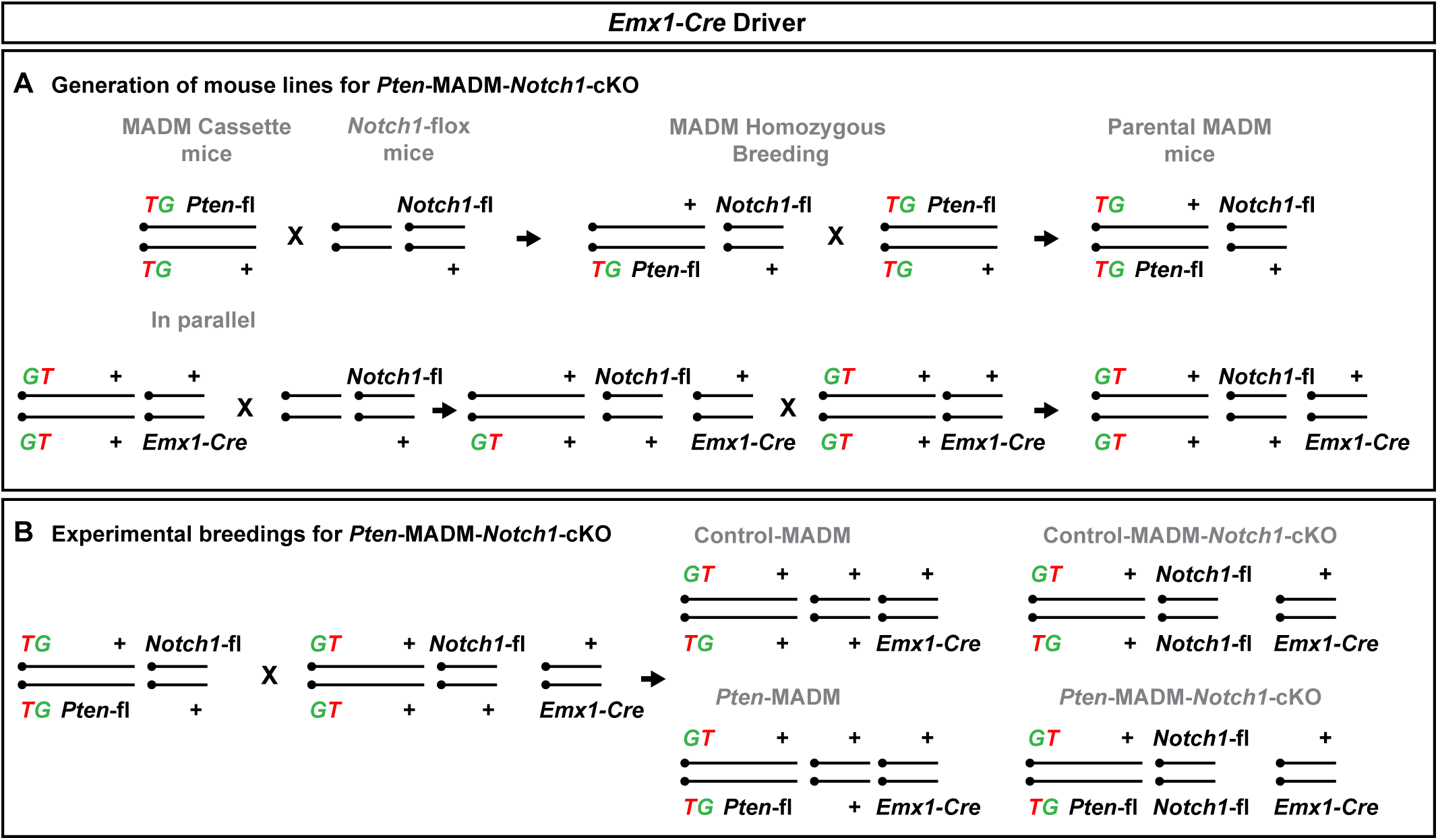
Breeding schemes to test for *Notch1* dependence in *Pten* mutant astrocyte phenotype. (A) Parental breeding scheme. *Notch1*-flox mice were crossed to *MADM-19^TG/TG,Pten^* and *MADM-19^GT/GT^*;*Emx1^Cre/+^*. The F_1_ offspring were heterozygous for the respective MADM cassette (*TG-Pten* or *GT*), *Emx1*-Cre, and *Notch1*-flox. These F_1_ offspring were crossed with *MADM-19^TG/TG,Pten^* and *MADM-19^GT/GT^*;*Emx1^Cre/+^*, respectively, to obtain MADM cassette homozygosity. The F_2_ offspring were used for strain maintenance and experimental breeding. (B) F_2_ offspring carrying MADM cassettes and *Notch1-*flox alleles were crossed to generate all four experimental genotypes as indicated.

**Figure S11.**
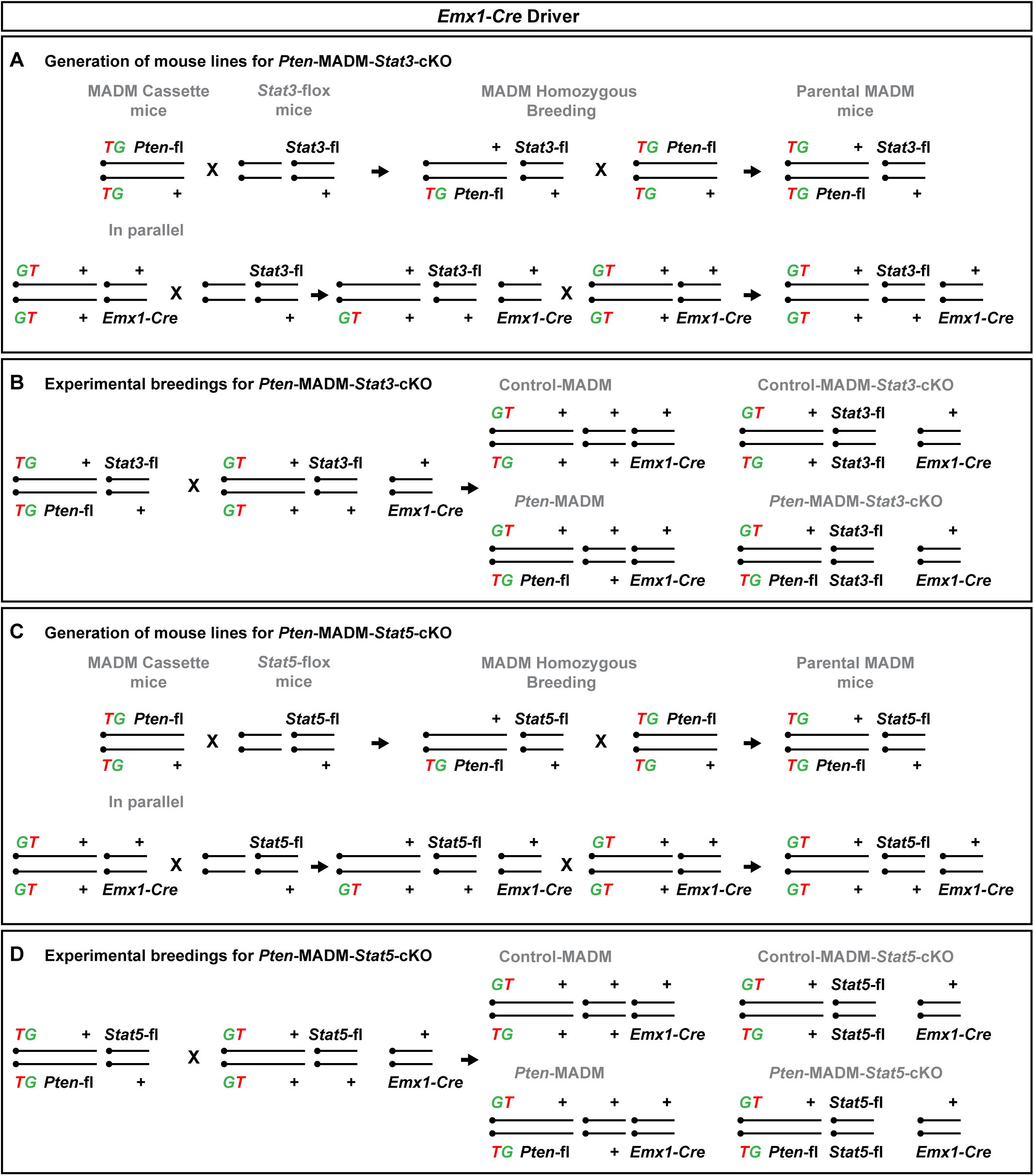
Breeding schemes to test for *Stat3* or *Stat5* dependence in *Pten* mutant astrocyte phenotype. (A) Parental breeding scheme. *Stat3*-flox mice were crossed to *MADM-19^TG/TG,Pten^* and *MADM-19^GT/GT^*;*Emx1^Cre/+^*. The F_1_ offspring were heterozygous for the respective MADM cassette (*TG-Pten* or *GT*), *Emx1*-Cre, and *Stat3*-flox. These F_1_ offspring were crossed with *MADM-19^TG/TG,Pten^* and *MADM-19^GT/GT^*;*Emx1^Cre/+^*, respectively, to obtain MADM cassette homozygosity. The F_2_ offspring were used for strain maintenance and experimental breeding. (B) F_2_ offspring carrying MADM cassettes and *Stat3-*flox alleles were crossed to generate all four experimental genotypes as indicated. (C) Parental breeding scheme. *Stat5*-flox mice were crossed to *MADM-19^TG/TG,Pten^* and *MADM-19^GT/GT^*;*Emx1^Cre/+^*. The F_1_ offspring were heterozygous for the respective MADM cassette (*TG-Pten* or *GT*), *Emx1*-Cre, and *Stat5*-flox. These F_1_ offspring were crossed with *MADM-19^TG/TG,Pten^* and *MADM-19^GT/GT^*;*Emx1^Cre/+^*, respectively, to obtain MADM cassette homozygosity. The F_2_ offspring were used for strain maintenance and experimental breeding. (D) F_2_ offspring carrying MADM cassettes and *Stat5-*flox alleles were crossed to generate all four experimental genotypes as indicated.

**Figure S12.**
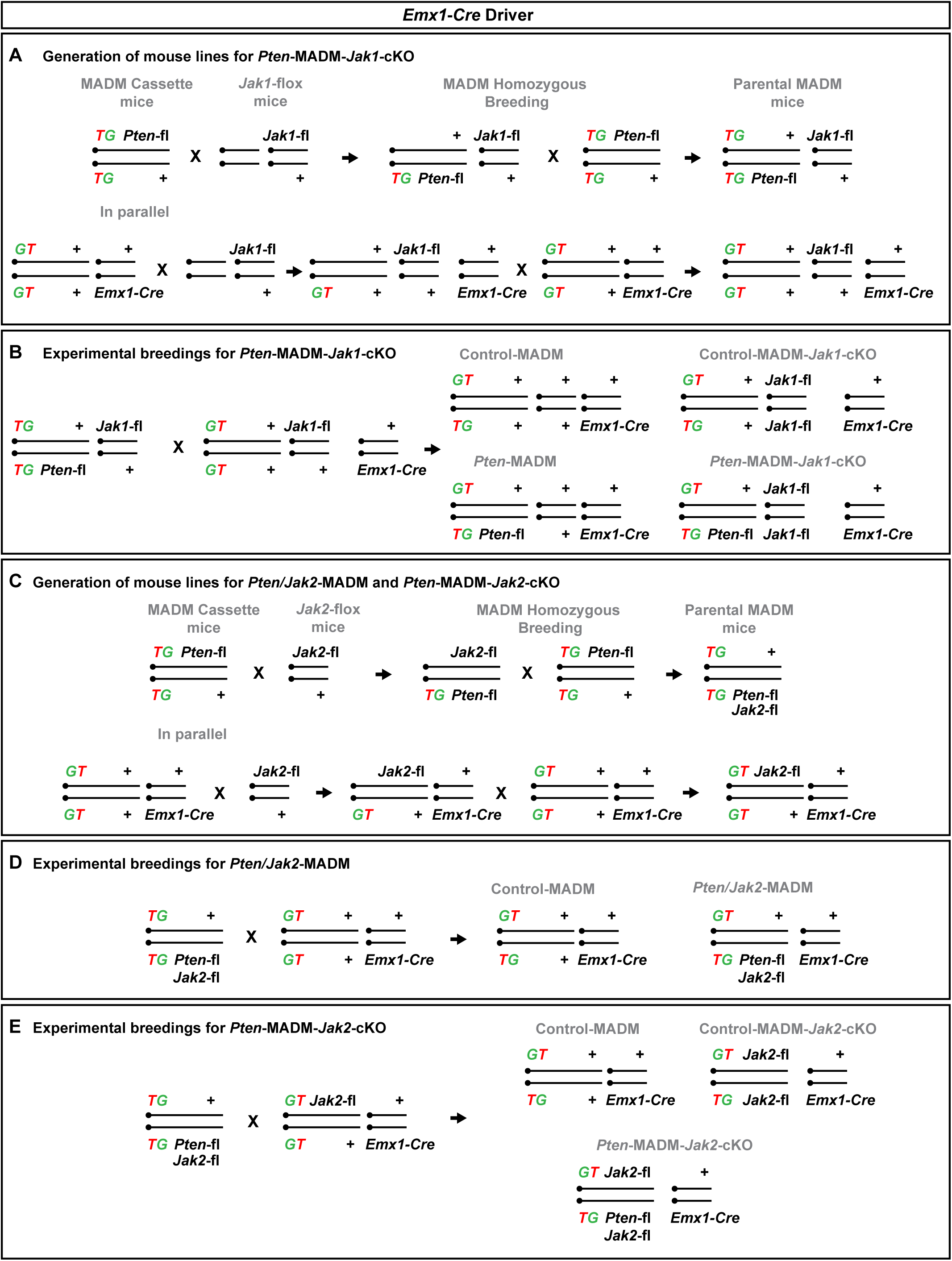
Breeding schemes to test for *Jak1* and/or *Jak2* dependence in *Pten* mutant astrocyte phenotype. (A) Parental breeding scheme. *Jak1*-flox mice were crossed to *MADM-19^TG/TG,Pten^* and *MADM-19^GT/GT^*;*Emx1^Cre/+^*. The F_1_ offspring were heterozygous for the respective MADM cassette (*TG-Pten* or *GT*), *Emx1*-Cre, and *Jak1*-flox. These F_1_ offspring were crossed with *MADM-19^TG/TG,Pten^* and *MADM-19^GT/GT^*;*Emx1^Cre/+^*, respectively, to obtain MADM cassette homozygosity. The F_2_ offspring were used for strain maintenance and experimental breeding. (B) F_2_ offspring carrying MADM cassettes and *Jak1-*flox alleles were crossed to generate all four experimental genotypes as indicated. (C) Parental breeding scheme. *Jak2*-flox mice were crossed to *MADM-19^TG/TG,Pten^* and *MADM-19^GT/GT^*;*Emx1^Cre/+^*. The F_1_ offspring were heterozygous for the respective MADM cassette (*TG-Pten* or *GT*), *Emx1*-Cre, and *Jak2*-flox. These F_1_ offspring were crossed with *MADM-19^TG/TG,Pten^* and *MADM-19^GT/GT^*;*Emx1^Cre/+^*, respectively, to obtain MADM cassette homozygosity, and *MADM-19^TG/TG,Jak2,Pten^* double recombinant. The F_2_ offspring were used for strain maintenance and experimental breeding. (D) F_2_ offspring carrying MADM cassettes and a double recombinant (i.e. TG-*Pten-*flox-*Jak2*-flox allele) were crossed to generate the double mosaic model (*Pten*-*Jak2*-MADM). Note: Since *Jak2* and *Pten* are both located on chr.19 (*Jak2* closer to centromere than *Pten*), it is statistically improbable to get a *Pten*-MADM animal (i.e. recombining *Jak2* away from *Pten*) in the same litter, hence we used Control-MADM in these comparisons. (E) F_2_ offspring carrying MADM cassettes and *Jak2-flox* alleles were crossed to generate all four experimental genotypes. Note: To generate the *Pten*-MADM-*Jak1/2*-cKO, the *Jak1*-flox allele was first crossed into the *MADM-19^TG/TG,Jak2,Pten^* double recombinant and *MADM-19^GT/GT,Jak2^*;*Emx1^Cre/+^*, respectively, to obtain *MADM-19^TG/TG,Jak2,Pten^*; *Jak1^flox/+^* and *MADM-19^GT/GT,Jak2^*;*Jak1^flox/+^*;*Emx1^Cre/+^*. These stocks were then used to generate the experimental *Pten*-MADM-*Jak1/2*-cKO (*MADM-19^GT,Jak2/TG,Jak2,Pten^*;*Jak1^flox/flox^*;*Emx1^Cre/+^*) animals.

**Figure S13.**
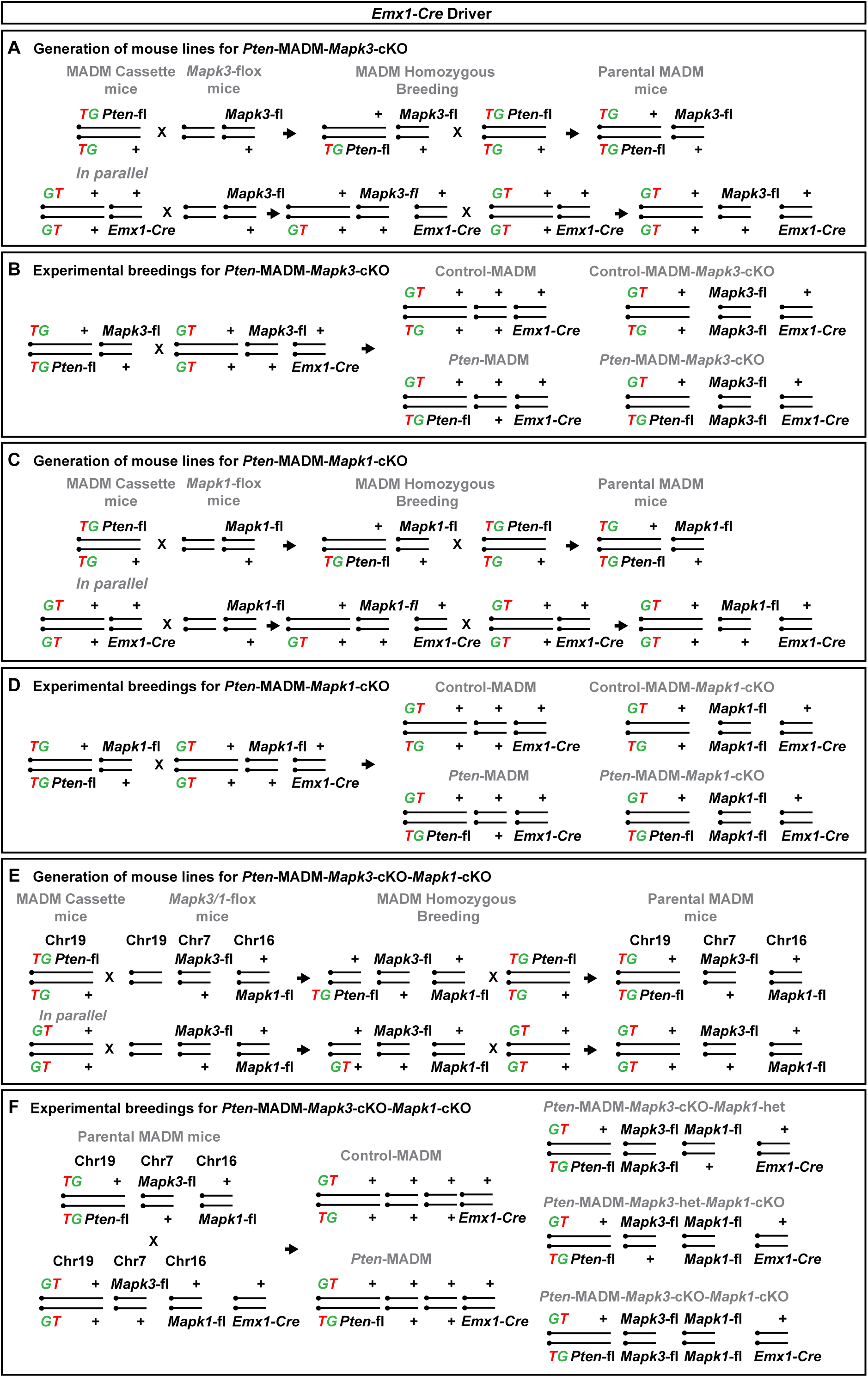
Breeding schemes to test for *Mapk3* and/or *Mapk1* dependence in *Pten* mutant astrocyte phenotype. (A) Parental breeding scheme. *Mapk3*-flox mice were crossed to *MADM-19^TG/TG,Pten^* and *MADM-19^GT/GT^*;*Emx1^Cre/+^*. The F_1_ offspring were heterozygous for the respective MADM cassette (*TG-Pten* or *GT*), *Emx1*-Cre, and *Mapk3*-flox. These F_1_ offspring were crossed with *MADM-19^TG/TG,Pten^* and *MADM-19^GT/GT^*;*Emx1^Cre/+^*, respectively, to obtain MADM cassette homozygosity. The F_2_ offspring were used for strain maintenance and experimental breeding. (B) F_2_ offspring carrying MADM cassettes and *Mapk3-*flox alleles were crossed to generate all four experimental genotypes as indicated. (C) Parental breeding scheme. *Mapk1*-flox mice were crossed to *MADM-19^TG/TG,Pten^* and *MADM-19^GT/GT^*;*Emx1^Cre/+^*. The F_1_ offspring were heterozygous for the respective MADM cassette (*TG-Pten* or *GT*), *Emx1*-Cre, and *Mapk1*-flox. These F_1_ offspring were crossed with *MADM-19^TG/TG,Pten^* and *MADM-19^GT/GT^*;*Emx1^Cre/+^*, respectively, to obtain MADM cassette homozygosity. The F_2_ offspring were used for strain maintenance and experimental breeding. (D) F_2_ offspring carrying MADM cassettes and *Mapk1-*flox alleles were crossed to generate all four experimental genotypes as indicated. (E) Parental breeding scheme. *Mapk1/3*-flox mice were crossed to *MADM-19^TG/TG,Pten^* and *MADM-19^GT/GT^*;*Emx1^Cre/+^*. The F_1_ offspring were heterozygous for the respective MADM cassette (*TG-Pten* or *GT*), *Emx1*-Cre, and *Mapk1/3*-flox. These F_1_ offspring were crossed with *MADM-19^TG/TG,Pten^* and *MADM-19^GT/GT^*;*Emx1^Cre/+^*, respectively, to obtain MADM cassette homozygosity. The F_2_ offspring were used for strain maintenance and experimental breeding. **(D)** F_2_ offspring carrying MADM cassettes and *Mapk1/3*-flox alleles were crossed to generate all five experimental genotypes as indicated.

## REFERENCES

Amberg N, Cheung G, Hippenmeyer S (2024) Protocol for sorting cells from mouse brains labeled with mosaic analysis with double markers by flow cytometry. STAR protocols 5 (1):102771. doi:10.1016/j.xpro.2023.102771

Amberg N, Pauler FM, Streicher C, Hippenmeyer S (2022) Tissue-wide genetic and cellular landscape shapes the execution of sequential PRC2 functions in neural stem cell lineage progression. Science advances 8 (44):eabq1263. doi:10.1126/sciadv.abq1263

An Z, Aksoy O, Zheng T, Fan QW, Weiss WA (2018) Epidermal growth factor receptor and EGFRvIII in glioblastoma: signaling pathways and targeted therapies. Oncogene 37 (12):1561–1575. doi:10.1038/s41388-017-0045-7

Asif M, Abdullah U, Nurnberg P, Tinschert S, Hussain MS (2023) Congenital Microcephaly: A Debate on Diagnostic Challenges and Etiological Paradigm of the Shift from Isolated/Non-Syndromic to Syndromic Microcephaly. Cells 12 (4). doi:10.3390/cells12040642

Backman SA, Stambolic V, Suzuki A, Haight J, Elia A, Pretorius J, Tsao MS, Shannon P, Bolon B, Ivy GO, Mak TW (2001) Deletion of Pten in mouse brain causes seizures, ataxia and defects in soma size resembling Lhermitte-Duclos disease. Nature genetics 29 (4):396–403. doi:10.1038/ng782

Balestrini S, Barba C, Thom M, Guerrini R (2023) Focal cortical dysplasia: a practical guide for neurologists. Practical neurology 23 (4):293–302. doi:10.1136/pn-2022-003404

Bandler RC, Vitali I, Delgado RN, Ho MC, Dvoretskova E, Ibarra Molinas JS, Frazel PW, Mohammadkhani M, Machold R, Maedler S, Liddelow SA, Nowakowski TJ, Fishell G, Mayer C (2022) Single-cell delineation of lineage and genetic identity in the mouse brain. Nature 601 (7893):404–409. doi:10.1038/s41586-021-04237-0

Barkovich AJ, Guerrini R, Kuzniecky RI, Jackson GD, Dobyns WB (2012) A developmental and genetic classification for malformations of cortical development: update 2012. Brain : a journal of neurology 135 (Pt 5):1348–1369. doi:10.1093/brain/aws019

Bartels T, Rowitch DH, Bayraktar OA (2024) Generation of Mammalian Astrocyte Functional Heterogeneity. Cold Spring Harbor perspectives in biology 16 (11). doi:10.1101/cshperspect.a041351

Beattie R, Postiglione MP, Burnett LE, Laukoter S, Streicher C, Pauler FM, Xiao G, Klezovitch O, Vasioukhin V, Ghashghaei TH, Hippenmeyer S (2017) Mosaic Analysis with Double Markers Reveals Distinct Sequential Functions of Lgl1 in Neural Stem Cells. Neuron 94 (3):517–533 e513. doi:10.1016/j.neuron.2017.04.012

Beattie R, Streicher C, Amberg N, Cheung G, Contreras X, Hansen AH, Hippenmeyer S (2020) Lineage Tracing and Clonal Analysis in Developing Cerebral Cortex Using Mosaic Analysis with Double Markers (MADM). Journal of visualized experiments : JoVE (159). doi:10.3791/61147

Bizzotto S, Walsh CA (2022) Genetic mosaicism in the human brain: from lineage tracing to neuropsychiatric disorders. Nature reviews Neuroscience 23 (5):275–286. doi:10.1038/s41583-022-00572-x

Blumcke I, Budday S, Poduri A, Lal D, Kobow K, Baulac S (2021) Neocortical development and epilepsy: insights from focal cortical dysplasia and brain tumours. The Lancet Neurology 20 (11):943–955. doi:10.1016/S1474-4422(21)00265-9

Brenner M, Kisseberth WC, Su Y, Besnard F, Messing A (1994) GFAP promoter directs astrocyte-specific expression in transgenic mice. The Journal of neuroscience : the official journal of the Society for Neuroscience 14 (3 Pt 1):1030–1037. doi:10.1523/JNEUROSCI.14-03-01030.1994

Burrows RC, Wancio D, Levitt P, Lillien L (1997) Response diversity and the timing of progenitor cell maturation are regulated by developmental changes in EGFR expression in the cortex. Neuron 19 (2):251–267. doi:10.1016/s0896-6273(00)80937-x

Cao J, Spielmann M, Qiu X, Huang X, Ibrahim DM, Hill AJ, Zhang F, Mundlos S, Christiansen L, Steemers FJ, Trapnell C, Shendure J (2019) The single-cell transcriptional landscape of mammalian organogenesis. Nature 566 (7745):496–502. doi:10.1038/s41586-019-0969-x

Casingal CR, Descant KD, Anton ES (2022) Coordinating cerebral cortical construction and connectivity: Unifying influence of radial progenitors. Neuron 110 (7):1100–1115. doi:10.1016/j.neuron.2022.01.034

Casingal CR, Nakagawa N, Yabuno-Nakagawa K, Meyer C, Liu S, Gkini V, Cho SJ, Skarica M, Liang D, Simon JM, Matoba N, Mallick A, Singla R, Park J, Huang CW, Wilson H, Lee J, Ghashghaei HT, Stuber GD, Heikinheimo O, Namba T, Stein JL, Anton ES (2025) TSC tunes progenitor balance and upper-layer neuron generation in neocortex. Nature. doi:10.1038/s41586-025-09810-5

Chen C, Ouyang W, Grigura V, Zhou Q, Carnes K, Lim H, Zhao GQ, Arber S, Kurpios N, Murphy TL, Cheng AM, Hassell JA, Chandrashekar V, Hofmann MC, Hess RA, Murphy KM (2005) ERM is required for transcriptional control of the spermatogonial stem cell niche. Nature 436 (7053):1030–1034. doi:10.1038/nature03894

Chen Y, Huang WC, Sejourne J, Clipperton-Allen AE, Page DT (2015) Pten Mutations Alter Brain Growth Trajectory and Allocation of Cell Types through Elevated beta-Catenin Signaling. The Journal of neuroscience : the official journal of the Society for Neuroscience 35 (28):10252–10267. doi:10.1523/JNEUROSCI.5272-14.2015

Chenn A, Walsh CA (2002) Regulation of cerebral cortical size by control of cell cycle exit in neural precursors. Science 297 (5580):365–369. doi:10.1126/science.1074192

Cheung G, Pauler FM, Koppensteiner P, Krausgruber T, Streicher C, Schrammel M, Gutmann-Ozgen N, Ivec AE, Bock C, Shigemoto R, Hippenmeyer S (2024) Multipotent progenitors instruct ontogeny of the superior colliculus. Neuron 112 (2):230–246 e211. doi:10.1016/j.neuron.2023.11.009

Chung WS, Baldwin KT, Allen NJ (2024) Astrocyte Regulation of Synapse Formation, Maturation, and Elimination. Cold Spring Harbor perspectives in biology 16 (8). doi:10.1101/cshperspect.a041352

Clark IC, Fontanez KM, Meltzer RH, Xue Y, Hayford C, May-Zhang A, D’Amato C, Osman A, Zhang JQ, Hettige P, Ishibashi JSA, Delley CL, Weisgerber DW, Replogle JM, Jost M, Phong KT, Kennedy VE, Peretz CAC, Kim EA, Song S, Karlon W, Weissman JS, Smith CC, Gartner ZJ, Abate AR (2023) Microfluidics-free single-cell genomics with templated emulsification. Nature biotechnology 41 (11):1557–1566. doi:10.1038/s41587-023-01685-z

Clavreul S, Abdeladim L, Hernandez-Garzon E, Niculescu D, Durand J, Ieng SH, Barry R, Bonvento G, Beaurepaire E, Livet J, Loulier K (2019) Cortical astrocytes develop in a plastic manner at both clonal and cellular levels. Nature communications 10 (1):4884. doi:10.1038/s41467-019-12791-5

Clavreul S, Dumas L, Loulier K (2022) Astrocyte development in the cerebral cortex: Complexity of their origin, genesis, and maturation. Frontiers in neuroscience 16:916055. doi:10.3389/fnins.2022.916055

Contreras X, Amberg N, Davaatseren A, Hansen AH, Sonntag J, Andersen L, Bernthaler T, Streicher C, Heger A, Johnson RL, Schwarz LA, Luo L, Rulicke T, Hippenmeyer S (2021) A genome-wide library of MADM mice for single-cell genetic mosaic analysis. Cell reports 35 (12):109274. doi:10.1016/j.celrep.2021.109274

Cui Y, Riedlinger G, Miyoshi K, Tang W, Li C, Deng CX, Robinson GW, Hennighausen L (2004) Inactivation of Stat5 in mouse mammary epithelium during pregnancy reveals distinct functions in cell proliferation, survival, and differentiation. Molecular and cellular biology 24 (18):8037–8047. doi:10.1128/MCB.24.18.8037-8047.2004

Currey L, Harvey T, Pelenyi A, Piper M, Thor S (2025) Mechanisms of brain overgrowth in autism spectrum disorder with macrocephaly. Frontiers in neuroscience 19:1586550. doi:10.3389/fnins.2025.1586550

Di Bella DJ, Dominguez-Iturza N, Brown JR, Arlotta P (2024) Making Ramon y Cajal proud: Development of cell identity and diversity in the cerebral cortex. Neuron 112 (13):2091–2111. doi:10.1016/j.neuron.2024.04.021

Di Cristofano A, Pesce B, Cordon-Cardo C, Pandolfi PP (1998) Pten is essential for embryonic development and tumour suppression. Nature genetics 19 (4):348–355. doi:10.1038/1235

Dobin A, Davis CA, Schlesinger F, Drenkow J, Zaleski C, Jha S, Batut P, Chaisson M, Gingeras TR (2013) STAR: ultrafast universal RNA-seq aligner. Bioinformatics 29 (1):15–21. doi:10.1093/bioinformatics/bts635

Fazekas F, Haboosheh A, Hennebichler B, Roetzer-Pejrimovsky T, Binder J, Reischer T, Pfeifer M, Scharrer A, Worda C, Linder T, Farr A, Hoftberger R, Gelpi E, Mitter C, Kasprian G, Haberler C, Amberg N (2026) A novel PTEN variant causing hemimegalencephaly and focal nodular heterotopias in the developing human brain. Epilepsia. doi:10.1002/epi.70088

Fenton TR, Nathanson D, Ponte de Albuquerque C, Kuga D, Iwanami A, Dang J, Yang H, Tanaka K, Oba-Shinjo SM, Uno M, Inda MM, Wykosky J, Bachoo RM, James CD, DePinho RA, Vandenberg SR, Zhou H, Marie SK, Mischel PS, Cavenee WK, Furnari FB (2012) Resistance to EGF receptor inhibitors in glioblastoma mediated by phosphorylation of the PTEN tumor suppressor at tyrosine 240. Proceedings of the National Academy of Sciences of the United States of America 109 (35):14164–14169. doi:10.1073/pnas.1211962109

Finak G, McDavid A, Yajima M, Deng J, Gersuk V, Shalek AK, Slichter CK, Miller HW, McElrath MJ, Prlic M, Linsley PS, Gottardo R (2015) MAST: a flexible statistical framework for assessing transcriptional changes and characterizing heterogeneity in single-cell RNA sequencing data. Genome biology 16:278. doi:10.1186/s13059-015-0844-5

Fraser MM, Bayazitov IT, Zakharenko SS, Baker SJ (2008) Phosphatase and tensin homolog, deleted on chromosome 10 deficiency in brain causes defects in synaptic structure, transmission and plasticity, and myelination abnormalities. Neuroscience 151 (2):476–488. doi:10.1016/j.neuroscience.2007.10.048

Fraser MM, Zhu X, Kwon CH, Uhlmann EJ, Gutmann DH, Baker SJ (2004) Pten loss causes hypertrophy and increased proliferation of astrocytes in vivo. Cancer research 64 (21):7773–7779. doi:10.1158/0008-5472.CAN-04-2487

Freeman MR, Rowitch DH (2013) Evolving concepts of gliogenesis: a look way back and ahead to the next 25 years. Neuron 80 (3):613–623. doi:10.1016/j.neuron.2013.10.034

Gao P, Postiglione MP, Krieger TG, Hernandez L, Wang C, Han Z, Streicher C, Papusheva E, Insolera R, Chugh K, Kodish O, Huang K, Simons BD, Luo L, Hippenmeyer S, Shi SH (2014) Deterministic progenitor behavior and unitary production of neurons in the neocortex. Cell 159 (4):775–788. doi:10.1016/j.cell.2014.10.027

Gao Y, Sun M, Fu T, Wang Z, Jiang X, Yang L, Liang XG, Liu G, Tian Y, Yang F, Li J, Li Z, Li X, You Y, Ding C, Wang Y, Ma T, Zhang Z, Xu Z, Chen B, Yang Z (2025) NOTCH, ERK, and SHH signaling respectively control the fate determination of cortical glia and olfactory bulb interneurons. Proceedings of the National Academy of Sciences of the United States of America 122 (9):e2416757122. doi:10.1073/pnas.2416757122

Ge WP, Miyawaki A, Gage FH, Jan YN, Jan LY (2012) Local generation of glia is a major astrocyte source in postnatal cortex. Nature 484 (7394):376–380. doi:10.1038/nature10959

Gorski JA, Talley T, Qiu M, Puelles L, Rubenstein JL, Jones KR (2002) Cortical excitatory neurons and glia, but not GABAergic neurons, are produced in the Emx1-expressing lineage. The Journal of neuroscience : the official journal of the Society for Neuroscience 22 (15):6309–6314. doi:10.1523/JNEUROSCI.22-15-06309.2002

Greig LC, Woodworth MB, Galazo MJ, Padmanabhan H, Macklis JD (2013) Molecular logic of neocortical projection neuron specification, development and diversity. Nature reviews Neuroscience 14 (11):755–769. doi:10.1038/nrn3586

Groszer M, Erickson R, Scripture-Adams DD, Lesche R, Trumpp A, Zack JA, Kornblum HI, Liu X, Wu H (2001) Negative regulation of neural stem/progenitor cell proliferation by the Pten tumor suppressor gene in vivo. Science 294 (5549):2186–2189. doi:10.1126/science.1065518

Gu Z, Hubschmann D (2023) simplifyEnrichment: A Bioconductor Package for Clustering and Visualizing Functional Enrichment Results. Genomics, proteomics & bioinformatics 21 (1):190–202. doi:10.1016/j.gpb.2022.04.008

Hagemann-Jensen M, Ziegenhain C, Chen P, Ramskold D, Hendriks GJ, Larsson AJM, Faridani OR, Sandberg R (2020) Single-cell RNA counting at allele and isoform resolution using Smart-seq3. Nature biotechnology 38 (6):708–714. doi:10.1038/s41587-020-0497-0

Hakanen J, Ruiz-Reig N, Tissir F (2019) Linking Cell Polarity to Cortical Development and Malformations. Frontiers in cellular neuroscience 13:244. doi:10.3389/fncel.2019.00244

Hanganu-Opatz IL, Butt SJB, Hippenmeyer S, De Marco Garcia NV, Cardin JA, Voytek B, Muotri AR (2021) The Logic of Developing Neocortical Circuits in Health and Disease. The Journal of neuroscience : the official journal of the Society for Neuroscience 41 (5):813–822. doi:10.1523/JNEUROSCI.1655-20.2020

Hansen AH, Duellberg C, Mieck C, Loose M, Hippenmeyer S (2017) Cell Polarity in Cerebral Cortex Development-Cellular Architecture Shaped by Biochemical Networks. Frontiers in cellular neuroscience 11:176. doi:10.3389/fncel.2017.00176

Hippenmeyer S (2023) Principles of neural stem cell lineage progression: Insights from developing cerebral cortex. Current opinion in neurobiology 79:102695. doi:10.1016/j.conb.2023.102695

Hippenmeyer S, Youn YH, Moon HM, Miyamichi K, Zong H, Wynshaw-Boris A, Luo L (2010) Genetic mosaic dissection of Lis1 and Ndel1 in neuronal migration. Neuron 68 (4):695–709. doi:10.1016/j.neuron.2010.09.027

Kessaris N, Fogarty M, Iannarelli P, Grist M, Wegner M, Richardson WD (2006) Competing waves of oligodendrocytes in the forebrain and postnatal elimination of an embryonic lineage. Nature neuroscience 9 (2):173–179. doi:10.1038/nn1620

Klingler E, Francis F, Jabaudon D, Cappello S (2021) Mapping the molecular and cellular complexity of cortical malformations. Science 371 (6527). doi:10.1126/science.aba4517

Krempler A, Qi Y, Triplett AA, Zhu J, Rui H, Wagner KU (2004) Generation of a conditional knockout allele for the Janus kinase 2 (Jak2) gene in mice. Genesis 40 (1):52–57. doi:10.1002/gene.20063

Kriegstein A, Alvarez-Buylla A (2009) The glial nature of embryonic and adult neural stem cells. Annual review of neuroscience 32:149–184. doi:10.1146/annurev.neuro.051508.135600

Kwon CH, Zhao D, Chen J, Alcantara S, Li Y, Burns DK, Mason RP, Lee EY, Wu H, Parada LF (2008) Pten haploinsufficiency accelerates formation of high-grade astrocytomas. Cancer research 68 (9):3286–3294. doi:10.1158/0008-5472.CAN-07-6867

Kwon CH, Zhu X, Zhang J, Knoop LL, Tharp R, Smeyne RJ, Eberhart CG, Burger PC, Baker SJ (2001) Pten regulates neuronal soma size: a mouse model of Lhermitte-Duclos disease. Nature genetics 29 (4):404–411. doi:10.1038/ng781

Lattke M, Guillemot F (2022) Understanding astrocyte differentiation: Clinical relevance, technical challenges, and new opportunities in the omics era. WIREs mechanisms of disease 14 (5):e1557. doi:10.1002/wsbm.1557

Laukoter S, Amberg N, Pauler FM, Hippenmeyer S (2020a) Generation and isolation of single cells from mouse brain with mosaic analysis with double markers-induced uniparental chromosome disomy. STAR protocols 1 (3):100215. doi:10.1016/j.xpro.2020.100215

Laukoter S, Pauler FM, Beattie R, Amberg N, Hansen AH, Streicher C, Penz T, Bock C, Hippenmeyer S (2020b) Cell-Type Specificity of Genomic Imprinting in Cerebral Cortex. Neuron 107 (6):1160–1179 e1169. doi:10.1016/j.neuron.2020.06.031

Lee TC, Threadgill DW (2009) Generation and validation of mice carrying a conditional allele of the epidermal growth factor receptor. Genesis 47 (2):85–92. doi:10.1002/dvg.20464

Lehtinen MK, Zappaterra MW, Chen X, Yang YJ, Hill AD, Lun M, Maynard T, Gonzalez D, Kim S, Ye P, D’Ercole AJ, Wong ET, LaMantia AS, Walsh CA (2011) The cerebrospinal fluid provides a proliferative niche for neural progenitor cells. Neuron 69 (5):893–905. doi:10.1016/j.neuron.2011.01.023

Li X, Newbern JM, Wu Y, Morgan-Smith M, Zhong J, Charron J, Snider WD (2012) MEK Is a Key Regulator of Gliogenesis in the Developing Brain. Neuron 75 (6):1035–1050. doi:10.1016/j.neuron.2012.08.031

Lin Y, Yang J, Shen Z, Ma J, Simons BD, Shi SH (2021) Behavior and lineage progression of neural progenitors in the mammalian cortex. Current opinion in neurobiology 66:144–157. doi:10.1016/j.conb.2020.10.017

Llorca A, Ciceri G, Beattie R, Wong FK, Diana G, Serafeimidou-Pouliou E, Fernandez-Otero M, Streicher C, Arnold SJ, Meyer M, Hippenmeyer S, Maravall M, Marin O (2019) A stochastic framework of neurogenesis underlies the assembly of neocortical cytoarchitecture. eLife 8. doi:10.7554/eLife.51381

Llorca A, Marin O (2021) Orchestrated freedom: new insights into cortical neurogenesis. Current opinion in neurobiology 66:48–56. doi:10.1016/j.conb.2020.09.004

Love MI, Huber W, Anders S (2014) Moderated estimation of fold change and dispersion for RNA-seq data with DESeq2. Genome biology 15 (12):550. doi:10.1186/s13059-014-0550-8

Ma Q, Chen G, Li Y, Guo Z, Zhang X (2024) The molecular genetics of PI3K/PTEN/AKT/mTOR pathway in the malformations of cortical development. Genes & diseases 11 (5):101021. doi:10.1016/j.gendis.2023.04.041

Ma Y, Sun S, Shang X, Keller ET, Chen M, Zhou X (2020) Integrative differential expression and gene set enrichment analysis using summary statistics for scRNA-seq studies. Nature communications 11 (1):1585. doi:10.1038/s41467-020-15298-6

Mahler M, Berard M, Feinstein R, Gallagher A, Illgen-Wilcke B, Pritchett-Corning K, Raspa M (2014) FELASA recommendations for the health monitoring of mouse, rat, hamster, guinea pig and rabbit colonies in breeding and experimental units. Laboratory animals 48 (3):178–192. doi:10.1177/0023677213516312

Markey KM, Saunders JC, Smuts J, von Reyn CR, Garcia ADR (2023) Astrocyte development-More questions than answers. Frontiers in cell and developmental biology 11:1063843. doi:10.3389/fcell.2023.1063843

Miranda OA, Cheung G, Hippenmeyer S (2024) Morphological Analysis of Neurons and Glia Using Mosaic Analysis with Double Markers. Methods in molecular biology 2831:283–299. doi:10.1007/978-1-0716-3969-6_19

Moh A, Iwamoto Y, Chai GX, Zhang SS, Kano A, Yang DD, Zhang W, Wang J, Jacoby JJ, Gao B, Flavell RA, Fu XY (2007) Role of STAT3 in liver regeneration: survival, DNA synthesis, inflammatory reaction and liver mass recovery. Laboratory investigation; a journal of technical methods and pathology 87 (10):1018–1028. doi:10.1038/labinvest.3700630

Molnar Z, Clowry GJ, Sestan N, Alzu’bi A, Bakken T, Hevner RF, Huppi PS, Kostovic I, Rakic P, Anton ES, Edwards D, Garcez P, Hoerder-Suabedissen A, Kriegstein A (2019) New insights into the development of the human cerebral cortex. Journal of anatomy 235 (3):432–451. doi:10.1111/joa.13055

Namba T, Huttner WB (2024) What Makes Us Human: Insights from the Evolution and Development of the Human Neocortex. Annual review of cell and developmental biology 40 (1):427–452. doi:10.1146/annurev-cellbio-112122-032521

Nowakowski TJ, Nano PR, Matho KS, Chen X, Corrigan EK, Ding W, Gao Y, Heffel M, Jayakumar J, Kaplan HS, Kronman FN, Kovner R, Mannens CCA, Song M, Steyert MR, Venkatesan S, Wallace JL, Wang L, Werner JM, Zhang D, Yuan G, Zuo G, Ament SA, Colantuoni C, Dulac C, Fan R, Gillis J, Kriegstein AR, Krienen FM, Kim Y, Linnarsson S, Mitra PP, Pollen AA, Sestan N, Tward DJ, van Velthoven CTJ, Yao Z, Bhaduri A, Zeng H (2025) The new frontier in understanding human and mammalian brain development. Nature 647 (8088):51–59. doi:10.1038/s41586-025-09652-1

Oliveira D, Leal GF, Sertie AL, Caires LC, Jr., Goulart E, Musso CM, Oliveira JRM, Krepischi ACV, Vianna-Morgante AM, Zatz M (2019) 10q23.31 microduplication encompassing PTEN decreases mTOR signalling activity and is associated with autosomal dominant primary microcephaly. Journal of medical genetics 56 (8):543–547. doi:10.1136/jmedgenet-2018-105471

Ortiz-Alvarez G, Daclin M, Shihavuddin A, Lansade P, Fortoul A, Faucourt M, Clavreul S, Lalioti ME, Taraviras S, Hippenmeyer S, Livet J, Meunier A, Genovesio A, Spassky N (2019) Adult Neural Stem Cells and Multiciliated Ependymal Cells Share a Common Lineage Regulated by the Geminin Family Members. Neuron 102 (1):159–172 e157. doi:10.1016/j.neuron.2019.01.051

Ozawa T, Riester M, Cheng YK, Huse JT, Squatrito M, Helmy K, Charles N, Michor F, Holland EC (2014) Most human non-GCIMP glioblastoma subtypes evolve from a common proneural-like precursor glioma. Cancer cell 26 (2):288–300. doi:10.1016/j.ccr.2014.06.005

Pirozzi F, Nelson B, Mirzaa G (2018) From microcephaly to megalencephaly: determinants of brain size. Dialogues in clinical neuroscience 20 (4):267–282. doi:10.31887/DCNS.2018.20.4/gmirzaa

Sakamoto K, Wehde BL, Radler PD, Triplett AA, Wagner KU (2016a) Generation of Janus kinase 1 (JAK1) conditional knockout mice. Genesis 54 (11):582–588. doi:10.1002/dvg.22982

Sakamoto K, Wehde BL, Yoo KH, Kim T, Rajbhandari N, Shin HY, Triplett AA, Radler PD, Schuler F, Villunger A, Kang K, Hennighausen L, Wagner KU (2016b) Janus Kinase 1 Is Essential for Inflammatory Cytokine Signaling and Mammary Gland Remodeling. Molecular and cellular biology 36 (11):1673–1690. doi:10.1128/MCB.00999-15

Samuels IS, Karlo JC, Faruzzi AN, Pickering K, Herrup K, Sweatt JD, Saitta SC, Landreth GE (2008) Deletion of ERK2 mitogen-activated protein kinase identifies its key roles in cortical neurogenesis and cognitive function. The Journal of neuroscience : the official journal of the Society for Neuroscience 28 (27):6983–6995. doi:10.1523/JNEUROSCI.0679-08.2008

Selcher JC, Nekrasova T, Paylor R, Landreth GE, Sweatt JD (2001) Mice lacking the ERK1 isoform of MAP kinase are unimpaired in emotional learning. Learning & memory 8 (1):11–19. doi:10.1101/lm.37001

Shannon P, Markiel A, Ozier O, Baliga NS, Wang JT, Ramage D, Amin N, Schwikowski B, Ideker T (2003) Cytoscape: a software environment for integrated models of biomolecular interaction networks. Genome research 13 (11):2498–2504. doi:10.1101/gr.1239303

Shen Z, Lin Y, Yang J, Jorg DJ, Peng Y, Zhang X, Xu Y, Hernandez L, Ma J, Simons BD, Shi SH (2021) Distinct progenitor behavior underlying neocortical gliogenesis related to tumorigenesis. Cell reports 34 (11):108853. doi:10.1016/j.celrep.2021.108853

Sibilia M, Steinbach JP, Stingl L, Aguzzi A, Wagner EF (1998) A strain-independent postnatal neurodegeneration in mice lacking the EGF receptor. The EMBO journal 17 (3):719–731. doi:10.1093/emboj/17.3.719

Skelton PD, Stan RV, Luikart BW (2020) The Role of PTEN in Neurodevelopment. Molecular neuropsychiatry 5 (Suppl 1):60–71. doi:10.1159/000504782

Song MS, Salmena L, Pandolfi PP (2012) The functions and regulation of the PTEN tumour suppressor. Nature reviews Molecular cell biology 13 (5):283–296. doi:10.1038/nrm3330

Stuart T, Butler A, Hoffman P, Hafemeister C, Papalexi E, Mauck WM, 3rd, Hao Y, Stoeckius M, Smibert P, Satija R (2019) Comprehensive Integration of Single-Cell Data. Cell 177 (7):1888–1902 e1821. doi:10.1016/j.cell.2019.05.031

Sullivan PF, Geschwind DH (2019) Defining the Genetic, Genomic, Cellular, and Diagnostic Architectures of Psychiatric Disorders. Cell 177 (1):162–183. doi:10.1016/j.cell.2019.01.015

Szklarczyk D, Kirsch R, Koutrouli M, Nastou K, Mehryary F, Hachilif R, Gable AL, Fang T, Doncheva NT, Pyysalo S, Bork P, Jensen LJ, von Mering C (2023) The STRING database in 2023: protein-protein association networks and functional enrichment analyses for any sequenced genome of interest. Nucleic acids research 51 (D1):D638–D646. doi:10.1093/nar/gkac1000

Taverna E, Gotz M, Huttner WB (2014) The cell biology of neurogenesis: toward an understanding of the development and evolution of the neocortex. Annual review of cell and developmental biology 30:465–502. doi:10.1146/annurev-cellbio-101011-155801

Tsang M, Dawid IB (2004) Promotion and attenuation of FGF signaling through the Ras-MAPK pathway. Science’s STKE : signal transduction knowledge environment 2004 (228):pe17. doi:10.1126/stke.2282004pe17

Veleva-Rotse BO, Barnes AP (2014) Brain patterning perturbations following PTEN loss. Frontiers in molecular neuroscience 7:35. doi:10.3389/fnmol.2014.00035

Villalba A, Gotz M, Borrell V (2021) The regulation of cortical neurogenesis. Current topics in developmental biology 142:1–66. doi:10.1016/bs.ctdb.2020.10.003

Wagner KU, Krempler A, Triplett AA, Qi Y, George NM, Zhu J, Rui H (2004) Impaired alveologenesis and maintenance of secretory mammary epithelial cells in Jak2 conditional knockout mice. Molecular and cellular biology 24 (12):5510–5520. doi:10.1128/MCB.24.12.5510-5520.2004

Williamson MR, Murali S, Deneen B (2025) Development and diversity of astrocytes. Development 152 (13). doi:10.1242/dev.204705

Worby CA, Dixon JE (2014) Pten. Annual review of biochemistry 83:641–669. doi:10.1146/annurev-biochem-082411-113907

Yang X, Klein R, Tian X, Cheng HT, Kopan R, Shen J (2004) Notch activation induces apoptosis in neural progenitor cells through a p53-dependent pathway. Developmental biology 269 (1):81–94. doi:10.1016/j.ydbio.2004.01.014

Yu G, Wang LG, Han Y, He QY (2012) clusterProfiler: an R package for comparing biological themes among gene clusters. Omics : a journal of integrative biology 16 (5):284–287. doi:10.1089/omi.2011.0118

Zhang X, Mennicke CV, Xiao G, Beattie R, Haider MA, Hippenmeyer S, Ghashghaei HT (2020) Clonal Analysis of Gliogenesis in the Cerebral Cortex Reveals Stochastic Expansion of Glia and Cell Autonomous Responses to Egfr Dosage. Cells 9 (12). doi:10.3390/cells9122662

Zhou J, Vitali I, Roig-Puiggros S, Javed A, Cantando I, Puglisi M, Bezzi P, Jabaudon D, Mayer C, Bocchi R (2025) Dual lineage origins contribute to neocortical astrocyte diversity. Nature communications 16 (1):6992. doi:10.1038/s41467-025-61829-4

Zong H, Espinosa JS, Su HH, Muzumdar MD, Luo L (2005) Mosaic analysis with double markers in mice. Cell 121 (3):479–492. doi:10.1016/j.cell.2005.02.012

